# Multiplex HDR for Disease and Correction Modeling of SCID by CRISPR Genome Editing in Human HSPCs

**DOI:** 10.1101/2022.10.07.511272

**Authors:** Ortal Iancu, Daniel Allen, Orli Knop, Yonathan Zehavi, Dor Breier, Adaya Arbiv, Atar Lev, Yu Nee Lee, Katia Beider, Arnon Nagler, Raz Somech, Ayal Hendel

**Affiliations:** The Institute for Advanced Materials and Nanotechnology, The Mina and Everard Goodman Faculty of Life Sciences, Bar-Ilan University, Ramat Gan, Israel; Pediatric Department A and the Immunology Service, Jeffrey Modell Foundation Center, Chaim Sheba Medical Center, Edmond and Lily Safra Children’s Hospital, Israel Ministry of Health, Tel-HaShomer, Ramat Gan, Israel; The Division of Hematology and Bone Marrow Transplantation, Chaim Sheba Medical Center, Tel-HaShomer, Ramat Gan, Israel; Sackler Faculty of Medicine, Tel Aviv University, Tel Aviv, Israel

## Abstract

Severe combined immunodeficiency (SCID) is a group of monogenic primary immunodeficiencies caused by mutations in genes involved in the process of lymphocyte maturation and function. CRISPR-Cas9 gene editing of the patient’s own hematopoietic stem and progenitor cells (HSPCs) *ex vivo* could provide a therapeutic alternative to allogeneic hematopoietic stem cell transplantation (HSCT), the current gold standard for treatment of SCID. Using CRISPR-Cas9/rAAV6 gene-editing, we engineered genotypes in healthy donor (HD)-derived CD34^+^ HSPCs, thus eliminating the need for rare patient samples, to model both SCID and the therapeutic outcomes of gene-editing therapies for SCID via multiplexed homology directed repair (HDR). Firstly, we developed a SCID disease model via *knock-out* of both alleles of genes critical to the development of lymphocytes; and secondly, we established a *knock-in/knock-out (KI-KO)* strategy to develop a proof-of-concept single-allelic gene correction. Since SCID is a recessive disorder, correction of only one allele is enough to cure the patient. Based on these results, we performed gene correction of *RAG2*-SCID patient-derived CD34^+^ HSPCs that successfully developed into CD3^+^ T cells with diverse TCR repertoires in an *in vitro* T-cell differentiation (IVTD) platform. By using CRISPR-Cas9, multiplexed HDR, HD-derived CD34^+^ HSPCs, and an IVTD system we outline an approach for the study of human lymphopoiesis. We present both a way for researchers to determine the optimal configuration for CRISPR-Cas9 gene correction of SCID and other recessive blood disorders, and the feasibility of translating these techniques to perform gene correction in patient-derived CD34^+^ HSPCs.

## Introduction

Severe combined immunodeficiency (SCID) is a group of multiple monogenic disorders characterized by a profound block of T-cell development which harms both cellular and humoral adaptive immunity^1,2^. Depending on the type of SCID, B and NK cells may also be affected. Thus, based on the affected gene, patients can be classified according to the presence or absence of T, B, and NK lymphocytes (T−/+, B−/+, and NK−/+, respectively). The most common SCID is X-linked SCID (SCID-X1 [from mutation in the *IL2RG* gene]), which presents with a T-B+NK-immune phenotype. Other forms of SCID can develop as a result of mutations in *Recombination-activating gene 1* (*RAG1*), *RAG2,* or *DNA Cross-Link Repair 1C* (*DCLRE1C*) and display the T-B-NK+ immune phenotype, while other forms develop from mutations in the *Interleukin 7 Receptor Subunit Alpha* (*IL7RA*) gene and present with a T-B+NK+ immune phenotype^3–7^. *RAG1* and *RAG2* genes encode proteins that, when complexed together, commence the lymphoid-specific variable (V), diversity (D), and joining (J) gene [V(D)J] recombination process by catalyzing DNA double-strand breaks (DSBs) at the recombination signal sequences (RSSs) which flank the V, D, and J gene segments^8^. The *DCLRE1C* gene encodes the Artemis protein which assists in the functional resection of the V, D, and J gene segments by employing its endonuclease activity on the 5’ and 3’ overhangs and hairpins of the RAG-complex-induced DSBs^9^. V(D)J recombination is a critical step in the maturation of T and B cells as it is responsible for the generation of a diverse repertoire of T and B cell receptors (TCR and BCR, respectively)^10^. Thus, patients with disease-causing variants in the *RAG1*, *RAG2*, or *DCLRE1C* genes typically present with significantly reduced or complete absence of T and B cells. IL7RA signaling plays a major role at various stages of T-cell development, namely ensuring the survival of naive T cells and assisting in the homeostatic expansion of both naive and memory T cells via proliferation^11^. Thus, typical patients with mutations in the *IL7RA* gene will present a lack of T cells. Ineffective expression of any of these genes can lead to SCID, highlighted by severe lymphopenia and lack of cellular and humoral adaptive immunity^6^.

Infants born with SCID appear healthy in the first few weeks of life, however, following environmental exposure to pathogens and the decline of maternally transferred antibodies, they become prone to develop life-threatening bacterial, viral, and/or fungal infections. Without early intervention to reconstitute their immune system, patients often do not survive past the first two years of life^2^. The definitive curable treatment for SCID patients is allogeneic HSCT^12^ from a human leukocyte antigen (HLA)-matched (related or unrelated) donor. If a full HLA-matched donor is not available, patients may undergo haploidentical HSCT from one of their parents. Successful HSCT promotes lymphoid lineage development resulting in a long-term patient survival rate of >80%, however, there are significant limitations to this approach especially in the absence of a HLA-matched donor where the survival rate decreases to 60-70%^13, 14^. These limitations include graft failure after HSCT that leads to poor immune reconstitution as well as potentially fatal graft-versus-host disease (GvHD)^15,16^. Therefore, due to the devastating nature of SCID and the limitations of HSCT, it is crucial to develop new treatment options, such as gene therapy.

The ability to genetically edit CD34^+^ hematopoietic stem and progenitor cells (HSPCs) as well as the cells’ marked ability to reconstitute the immune system from a small number of original cells, make these stem cells an attractive target for gene therapy^17,18^. Viral vectors can facilitate the delivery of a corrected transgene to autologous HSPCs *ex vivo*^19,20^ as was previously established in gene therapy clinical trials using lentiviral (LV) or gammaretroviral (γRV) transduction^21–23^. Treatment of Chronic Granulomatous Disease (CGD)^24^, Wiskott-Aldrich Syndrome (WAS)^25,26^, SCID-X1^24,27^, and most recently of Adenosine Deaminase (ADA)-SCID with γRV resulted in the activation of proto-oncogenes leading to a leukemic transformation in some patients^28,29^. To improve the safety of viral transgene delivery, LVs have replaced γRVs, the viral enhancer sequences responsible for the elevated risk of genotoxicity were removed^30^, and a self-inactivating element was added to the vectors. This resulted in successful therapy for SCID-X1 patients, however, expansion of a single clonal population was reported in one patient case^31–34^. The main disadvantages of using γRV and LV vectors for CD34^+^ HSPC gene therapy remain the semi-random integration and constitutive expression of the transgene which can lead to incomplete phenotypic correction, dysregulated hematopoiesis, toxicity, and insertional mutagenesis^35^. Thus, delivery of the transgene via a targeted genome-editing approach could prove beneficial.

Clustered regularly interspaced short palindromic repeats [CRISPR] and CRISPR-associated nuclease 9 [Cas9], commonly known as CRISPR-Cas9, has had a tremendous impact on the field of gene editing due to its simplicity, specificity, and applicability in a wide variety of cell types^36–38^. The combination of CRISPR-Cas9 and a donor DNA molecule delivered by recombinant adeno-associated virus serotype 6 (rAAV6) can provide a therapeutic approach to genome editing in CD34^+^ HSPCs. Although rAAV6 lacks any integration machinery of its own^39^, precise targeting and insertion of the donor DNA payload through the homology-directed repair (HDR) pathway in CD34^+^ HSPCs can occur following the Cas9-induced site-specific DSB^40,41^. However, use of rAAV6 vectors is not without challenges. Our group has shown that rAAV6 vectors are recognized by cellular repair proteins triggering a DNA damage response (DDR) proportional to that of the amount of virus used, referred to as the multiplicity of infection (MOI)^42^. Therefore, when developing rAAV6-based protocols, reducing the MOI as much as possible, while maintaining the required levels of HDR, is of utmost importance.

Developing a gene-therapy strategy requires a thorough understanding of the disease phenotype and extensive assessment of the viability of the respective functional gene correction before it can reach the clinic. To achieve this, large amounts of patient samples are required to model and test the method *ex vivo*. Since SCID patients are infants, neither drawing large amounts of peripheral blood (PB) nor retrieving large samples through invasive bone-marrow procedures are viable options for procuring such samples. Therefore, obtaining sufficient amounts of SCID patient-derived CD34^+^ HSPCs presents a major challenge. To circumvent the need for large amounts of patient samples, we utilized multiplex HDR, which has been shown to be an effective method for enrichment of cells with engineered genotypes^43^. We engineered HD-derived CD34^+^ HSPCs to 1) model SCID and 2) simulate gene-correction therapies for SCID. Via cell sorting we were then able to enrich for cells with the desired engineered genotype to model and track their T-cell progression.

In this study, we accomplished three main goals towards bringing a curative gene therapy for SCID closer to becoming a reality, while providing a more general technique for modeling other recessive blood disorders using HD-derived CD34^+^ HSPCs. (1) We developed an innovative autosomal recessive SCID disease model via biallelic *knock-out* (*KO*) of *RAG2*, *DCLRE1C,* or *IL7RA* genes in HD-derived CD34^+^ HSPCs. When *KO* CD34^+^ HSPCs were subjected to a stromal cell-free, IVTD system^44^, these engineered cells lacked the ability to differentiate into CD3-expressing cells and presented a failure in execution of TCR V(D)J recombination, similar to that of SCID-patient-derived cells. (2) We then utilized a *KI-KO* approach to simulate functional gene correction of *RAG2* in HD-derived CD34^+^ HSPCs. Due to the recessive nature of SCID, correction of only one allele is sufficient to cure the patient. In this strategy, we mimicked monoallelic correction in SCID-patient cells by *knock-in* (*KI*) of a codon-optimized diverged cDNA cassette into the endogenous *RAG2* loci in one allele (thereby preserving regulatory non-coding elements) and *KO* of the second allele. In contrast to the *RAG2* biallelic KO cells, the *KI-KO* differentiated T cells presented normal CD3 expression and TCR repertoire diversity. (3) Lastly, we showed first-of-its-kind functional gene correction of *RAG2*-SCID patient-derived CD34^+^ HSPCs. In contrast to the unedited *RAG2*-SCID-patient cells, the corrected population developed into CD3^+^ T cells with diverse TCR repertoires. Overall, our method provides a platform to study the disease phenotypes with a multi-parameter readout in the form of immunophenotyping and V(D)J recombination assessment via next-generation sequencing (NGS) and will allow researchers to determine the optimal configuration for gene therapies for other SCIDs and immunodeficiencies.

## Results

### Modeling SCID disease with CRISPR-Cas9/rAAV6-mediated biallelic *KO* in HD-derived CD34^+^ HSPCs

In order to generate biallelic *KO* of the *RAG2*, *DCLRE1C,* or *IL7RA* genes individually in HD-derived CD34^+^ HPSCs, chemically modified sgRNAs^37^ were designed to target the genomic DNA a few base-pairs downstream of the respective gene’s start codon. The sgRNA was delivered together with the Cas9 endonuclease as a ribonucleoprotein (RNP) complex in conjunction with rAAV6 vectors carrying a DNA template for gene disruption. The template contained a reporter gene flanked by arms of homology for HDR at the aforementioned CRISPR-Cas9 cut-site. Following successful HDR, the integration of the reporter gene into the target gene’s open reading frame was expected to abolish the transcription of the coding region of the target gene (Figure 1A). Two DNA donors, each containing a distinct reporter gene, were required for each gene to allow for multiplex HDR and biallelic *KO* enrichment via cell sorting of cells expressing both reporter genes. We used two control groups in the following analyses: (1) cells electroporated without the presence of the RNP complex or rAAV6 (herein referred to as *Mock*) as well as (2) cells with biallelic *KO* of the *C-C motif chemokine receptor 5* (*CCR5*) gene. While *CCR5* is expressed in T cells, its expression does not affect T-cell development, allowing for the determination of the effect of CRISPR-Cas9 and rAAV6 treatments on the T-cell developmental process. Biallelic targeting of *RAG2* and *CCR5* were carried out with green fluorescent protein (GFP) and truncated nerve growth factor receptor (tNGFR) reporter genes (Figure 1A and S1A), whereas biallelic targeting of *DCLRE1C* and *IL7RA* were carried out with GFP and tdTomato reporter genes (Figure S1B and S1C). Immediately following electroporation, cells were exposed to their respective rAAV6 donors. Two days post-electroporation, the cells were sorted for CD34 expression, and double-positive reporter gene expression tNGFR^+^/GFP^+^ or tdTomato^+^/GFP^+^, indicative of biallelic *KO* (See example for *RAG2* in Figure 1B). Biallelic HDR frequencies at the different loci ranged from 1.7-4.6% (Figure 1C and S1D). DNA was purified from cells that were cultured in CD34^+^ HSPCs medium post-sort and the DNA was analyzed by Digital Droplet PCR (ddPCR) for quantification of locus-specific target integration. ddPCR analysis revealed that the *CCR5, RAG2*, and *DCLRE1C* sorted cells contained the *KO* disruption DNA donors in all targeted alleles whereas *IL7RA* sorted cells showed slightly less efficient integration (Figure S1E).

**Figure 1.**
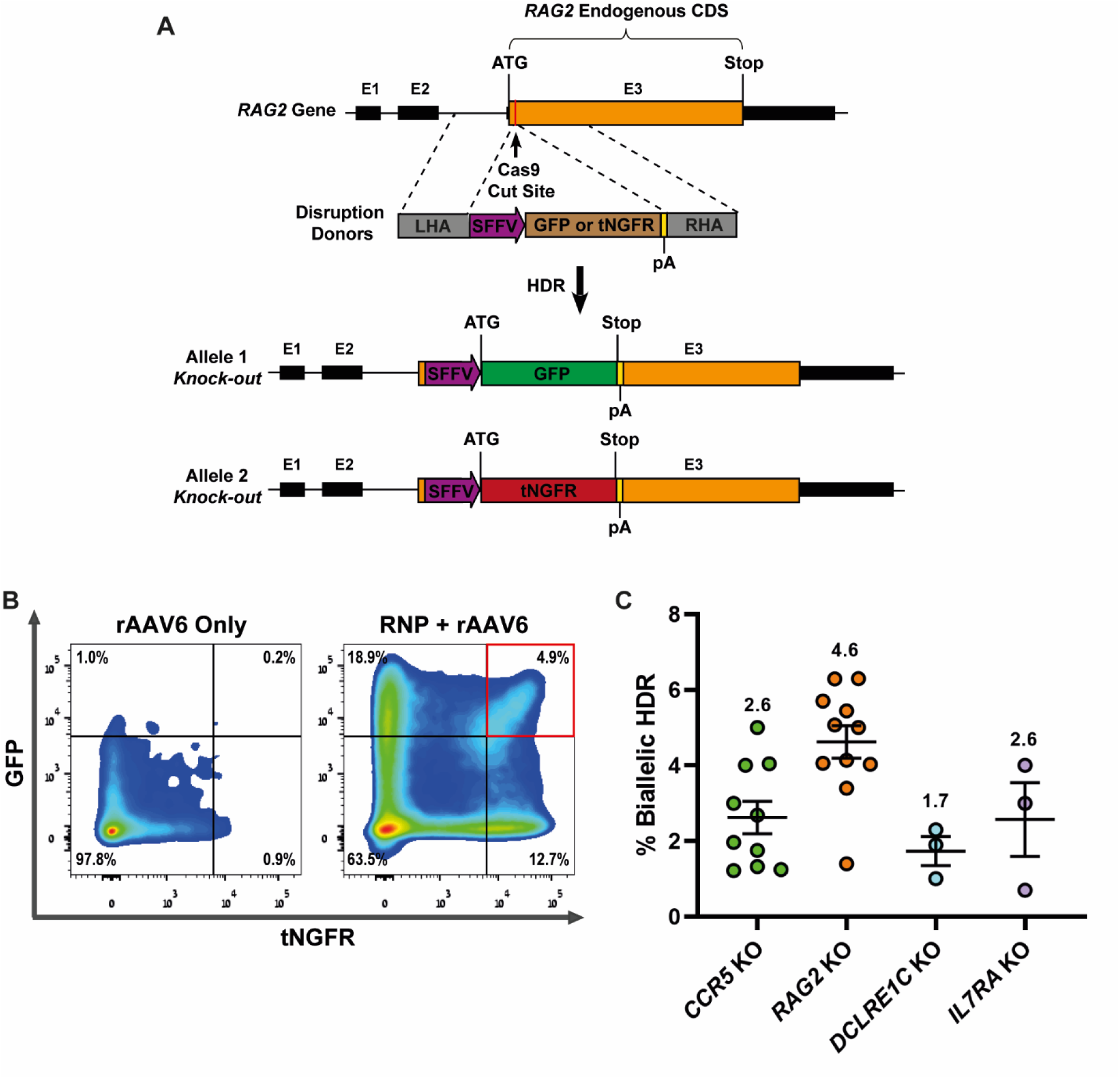
Biallelic targeting of SCID-related genes via CRISPR-Cas9/rAAV6 in HD-derived CD34^+^ HSPCs. **(A)** Schematic representation of *RAG2* disruption donors containing tNGFR or GFP selectable markers driven by an SFFV promoter and two 400 bp arms of homology for direct recombination repair (left homologous arm [LHA]; right homologous arm [RHA]). Successful HDR results in *KI* of the reporter gene approximately 43 bp downstream from the *RAG2* ATG start codon. *DCLRE1C* and *IL7RA* biallelic editing (not shown here) utilize tdTomato and GFP reporter genes. **(B)** FACS approach for enrichment of biallelic *RAG2* gene-targeted CD34^+^ HSPCs 2 days after CRISPR-Cas9 and rAAV6 editing. Representative HSPC FACS plots of cells transduced with rAAV6 only (*left*) and cells treated with CRISPR-Cas9 and rAAV6 (*right*) are shown. The double-positive tNGFR^+^/GFP^+^ population indicative of biallelic integration of two different DNA donors were sorted. Gating determination is based on cells treated with only the rAAV6 vector (left) to compensate for the episomal expression determined to be around 1%. **(C)** GFP and either tNGFR or tdTomato biallelic targeting frequencies at *CCR5* (N=10), *RAG2* (N=11), *DCLRE1C* (N=3), and *IL7RA* loci (N=3). Data are represented as mean ± SEM.

### Modeled SCID CD34^+^ HSPCs do not progress to CD3^+^ T Cells in a cell-free *in vitro* T-cell differentiation system

CD34^+^ HSPCs differentiate into CD7^+^/CD5^+^ pro-T cells which then continue to develop into T-cell-committed pre-T CD1a^+^ cells (identifiable in IVTD by immunostaining on day 14). These CD7^+^/CD1a^+^ cells can then undergo TCR rearrangement and become immature single-positive (ISP) CD4^+^ T cells. These give rise to double-positive (DP) CD4^+^/CD8^+^ cells that express CD3 (identifiable in IVTD by immunostaining on day 28 and 42 [in cases where extensive follow-up is required]). These cells then undergo full maturation into CD3-expressing single-positive (SP) CD8^+^ or CD4^+^ cells (i.e. CD3^+^/CD8^+^ or CD3^+^/CD4^+^ cells [Figure S2A]). CD3 receptors form a complex with the cell’s TCR, which serves a fundamental role in the maturation of thymocytes from their immature precursors. Typical SCID is characterized by the absence of or significant reduction in CD3^+^ T cells (<300 cells/ml)^45^. The stromal cell-free IVTD system that we used provided the ability for easy observation and immunophenotyping by flow cytometry of T-cell development at predetermined time points and assessment of V(D)J recombination via NGS analysis.

In order to model *RAG2-, DCLRE1C-,* or *IL7RA*-SCID, *biallelic KO* CD34^+^ HSPCs for each locus were individually cultured in the IVTD system and compared to *Mock* and *CCR5* KO populations. Consistent with the literature, the *IL7RA* biallelic KO populations presented complete cell death after only 6 days in culture^6^ (Figure 2A), whereas cells that had no HDR integration were predominantly viable cells (Figure S2B). In contrast, *Mock* cells and *CCR5*, *RAG2*, and *DCLRE1C* biallelic KO populations were able to proliferate through the 28 day time point (Figure 2B). Immunophenotyping by flow cytometry revealed no significant difference in the expression levels of CD5, CD7, or CD1a between the *RAG2* or *DCLRE1C* KO and *CCR5* KO cells after 14 days. Additionally, similar expression of both CD4 and CD8 were observed in all three groups after 28 days. However, at this stage of differentiation, *CCR5* KO cells developed into CD3^+^ T cells, while hardly any *RAG2* or *DCLRE1C* KO cells expressed CD3 (Figure 2B and S2C). Furthermore, as expected, the expression of the endogenous *RAG2* and *DCLRE1C* genes were drastically reduced in the *RAG2* and *DCLRE1C* biallelic *KO* populations (Figure S2D-E).

**Figure 2.**
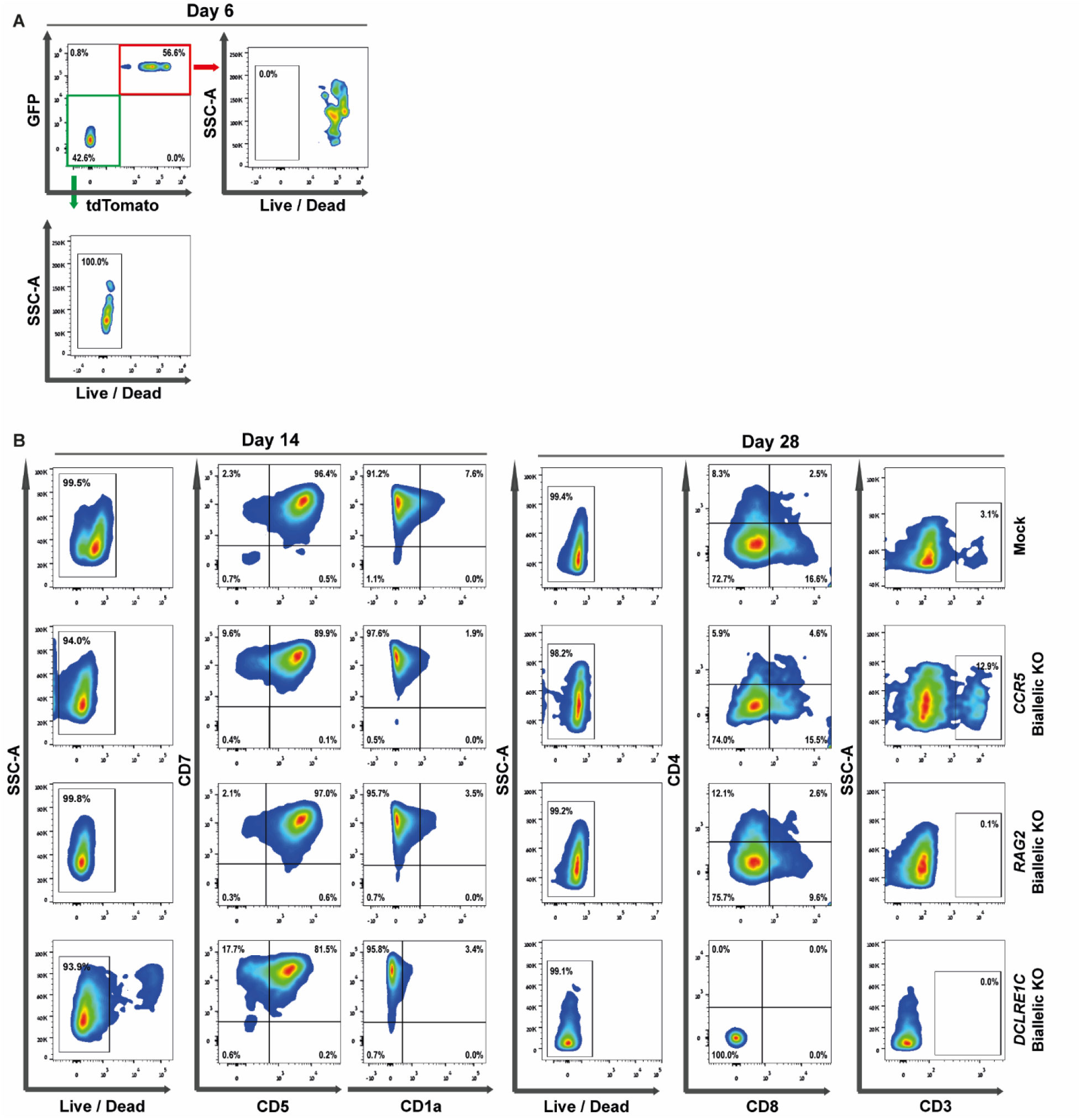
SCID modeling using biallelic *KO* of HD-derived CD34^+^ HSPCs in a cell-free IVTD assay. **(A)** Flow cytometry analysis of *IL7RA* biallelic KO in IVTD. *IL7RA* biallelic KO cells showed no survival by day 6 of IVTD, based on flow cytometry viability stain out of the double-positive tdTomato^+^/GFP^+^ biallelic enriched population. Double-negative (tdTomato^−^/GFP^−^) cells showed 100% survival on day 6. Gating was determined by unstained cells. The data represent one of 3 independent experiments. **(B)** Flow cytometry analysis of T-cell developmental progression. *Mock*, *CCR5* biallelic KO, *RAG2* biallelic KO, and *DCLRE1C* biallelic KO cells developed normal expression of early markers of T-cell differentiation (CD7, CD5, and CD1a) upon analysis at 14 days of IVTD. Following 28 days of IVTD, *Mock* and *CCR5* biallelic KO cells expressed mature T-cell markers: CD4, CD8, and CD3, while *RAG2* biallelic KO and *DCLRE1C* biallelic KO cells lack CD3 expression. Gating was determined by FMO + isotype controls. The data represent one of 3-10 independent repeats (*Mock* [day 14 N=10 and day 28 N=7], *CCR5* KO [day 14 N=10 and day 28 N=6], *RAG2* KO [day 14 N=10 and day 28 N=6], and *DCLRE1C* KO [day 14 N=3 and day 28 N=3]). Cell viability for all samples remained >98% through day 28.

Due to their central role in somatic recombination, we sought to determine whether the *RAG2* and *DCLRE1C* KO cells were able to perform functional V(D)J recombination, the lack of which is a hallmark of SCID lymphocytes. To do so, we deep-sequenced the T-cell receptor gamma (TRG) loci, one of the first chains to be recombined among the different TCR chains^46,47^. Preparation of TRG libraries from *RAG2* KO cells revealed no amplification of the recombined locus (Figure S2F) and the rearrangement in *DCLRE1C* KO cells was similarly impaired, noted by standard PCR amplification using primers flanking the V-J regions of the TRG locus (Figure S2G). Taken together, our IVTD approach can be used to accurately model SCID and help us characterize the SCID phenotype for validation of future gene-correction results. This was highlighted by accurately producing the gene-specific disease phenotypes: *IL7RA* KO cells did not proliferate or differentiate at all and *DCLRE1C* KO and *RAG2* KO cells neither developed into CD3^+^ T cells nor underwent effective TCR V(D)J recombination.

### T-cell differentiation phenotype of SCID-patient cells validates the disease model

To validate our use of the IVTD platform as a reliable method to model the SCID phenotype, we subjected SCID-patient-derived CD34^+^ HSPCs to the system and examined their differentiation capability (Figure S3A). The SCID-patient-derived CD34^+^ HSPCs samples were extracted and purified from the PB of *RAG1*- and *RAG2*-SCID patients (Figure S3B-C). Both patients presented with a clinical phenotype suggestive of SCID and subsequent immune workup and genetic testing confirmed the diagnosis. The first patient is homozygous for a 4bp deletion (c.1407-10 del.4bp-TTGC) in the *RAG1* gene, resulting in a frameshift mutation, premature stop codon, and dysfunctional RAG1 protein^8^, while the second patient is homozygous for a missense mutation (c.G104T; p.G35V^48^) in the *RAG2* gene, which reduces the binding capabilities of RAG2 to RAG1 during V(D)J recombination. Alongside the CD34^+^ HSPCs from each patient, control HD-derived PB CD34^+^ HSPCs were seeded in the IVTD system.

After 14 days, there was no significant immunophenotypic difference in CD5 or CD7 expression between the groups, however, a range of CD1a expression was noticed across the sample populations (0-37%) (Figure S3A). Following 28 and 42 days in the IVTD system, the SCID-patient-derived cells did not differentiate into CD3^+^ cells and cell viability was still >90% in all samples, indicative of a halt in T-cell progression (Figure S3A). To emphasize the V(D)J recombination impairment in the SCID-patient cells compared to the HD-derived PB control, DNA was extracted and used as a template for TRG recombination PCR. The HD-derived PB cells successfully underwent TRG recombination, whereas the SCID-patient-derived PB cells failed (Figure S3D-F). An additional sample of cells was extracted and purified from PB of the *RAG1*-SCID patient following successful HSCT and subjected to the IVTD system. Consistent with the clinical reports, the *RAG1*-SCID cells post-HSCT showed robust CD3 expression (Figure S3A and Table S1) and completed successful TRG V(D)J recombination by day 14 (Figure S3F).

### Biallelic *KI-KO* targeting of *RAG2* in HD-derived CD34^+^ HSPCs to simulate functional gene correction

Since *RAG2*-SCID is a recessive disorder, correction of only one allele is required to have the patient develop a functional immune system. This is highlighted by the fact that carriers of *RAG2* mutations (e.g. the parents of the *RAG2*-SCID patient) present as fully healthy individuals. We aimed to utilize multiplex HDR for *RAG2 KI-KO* for correction simulation in HD-derived CD34^+^ HSPCs, which provided three main advantages over alternative editing strategies such as first *RAG2 KO* and then correction of the previously KO cells. 1) In contrast to prior works that have used induced pluripotent stem cells (iPSCs), HD-derived CD34^+^ HSPCs provided us with the capability of using biologically-authentic CD34^+^ HSPCs, the same cells that are used in HSCT^49^; 2) Culturing CD34^+^ HSPCs for too long reduces the cells regenerative potential. CD34^+^ HSPCs maintain their stemness and a balance between proliferation, quiescence, and regeneration as well as their ability to differentiate to all hematopoietic cell lineages for a limited time in culture. Extensive culturing protocols with multiple editing steps have been shown to lead to a loss of the stem cells qualities and engraftment potential in murine models^50^; and 3) It provides the ability to circumvent the need for rare patient samples to establish the feasibility of our correction strategy.

Based on this, we aimed to simulate functional gene correction of *RAG2* in HD-derived CD34^+^ HSPCs via multiplex HDR using two distinct rAAV6 donors to actuate biallelic *KI-KO*. In the HD-derived CD34^+^ HSPCs, we targeted one allele with a *KI* template containing diverged functional *RAG2* cDNA (along with a [tNGFR] reporter gene cassette under the regulation of a constitutive phosphoglycerokinase (PGK) promoter) (Figure 3A) and the other allele with a *KO* template consisting of a disrupting reporter cassette (*KO* schematic depicted in *Figure 1A* [GFP reporter gene under the regulation of a constitutive spleen focus-forming virus {SFFV} promoter.]). In this context, the use of the tNGFR reporter is advantageous since it enables tracking and enrichment of the corrected cells and has been approved for clinical applications^51^. Our *KI-KO* approach mimics the correction in recessive SCID patient cells where a single functional allele is enough to confer the host with a normal immune system. The *KI* correction donor DNA contained complete *RAG2* cDNA with silent mutations added to diverge the cDNA sequence from that of endogenous while maintaining codon usage. The resulting diverged cDNA produces the correct protein sequence, while the reduced similarity to the genomic sequence precludes the cDNA sequence from being re-cut by residual Cas9 or from serving as a potential homology arm causing premature cessation of HDR^52^. These *KI-KO* cells, double-positive for GFP and tNGFR, simulate the genotype and therapeutic outcome of *RAG2*-SCID single-allelic correction gene-editing therapy and were analyzed relative to our disease modeled biallelic *RAG2* KO cells.

**Figure 3.**
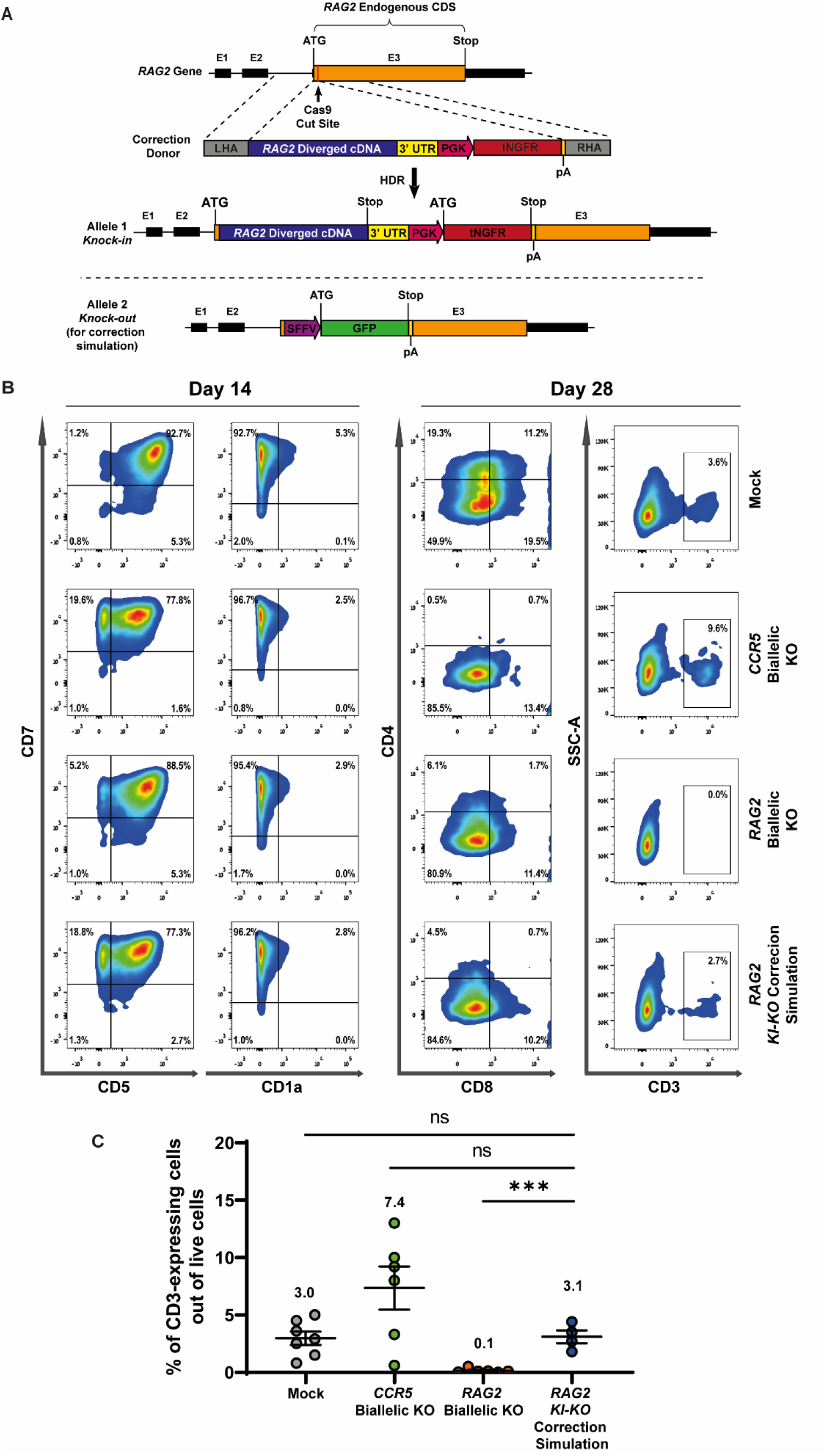
*RAG2 KI-KO* gene correction simulation strategy for in HD-derived CD34^+^ HSPCs. **(A)** Schematic representation of *RAG2* correction donor containing diverged codon-optimized *RAG2* cDNA and 3’ UTR followed by a tNGFR selectable marker driven by a PGK promoter. This is flanked by two 400 bp arms of homology for direct recombination repair, labeled LHA and RHA, on the left and right side of the CRISPR cut site, respectively. **(B)** Flow cytometry analysis of T-cell developmental progression of *Mock*, *CCR5* biallelic KO, *RAG2* biallelic KO, and *RAG2 KI-KO* correction simulation cells. All of the groups showed normal expression of early markers: CD7, CD5, and CD1a (upon 14 days of IVTD), as well as mature T-cell markers: CD4 and CD8 (upon 28 days of IVTD). CD3 expression was observed in *Mock*, *CCR5* biallelic KO, and *RAG2 KI-KO* correction simulation cells, yet not in *RAG2* biallelic KO cells. **(C)** Summary of CD3 expression by *Mock*, *CCR5* biallelic KO, *RAG2* biallelic KO, and *RAG2 KI-KO* correction simulation populations at day 28 of IVTD (N=7, N=6, N=6, and N=4, respectively). The data for *Mock*, *CCR5* KO, and *RAG2* KO are taken from *Figure S2C* and are presented here for comparison to the *RAG2 KI-KO* correction simulation population. *** p<0.0005 (*t*-test). Data are represented as mean ± SEM.

*RAG2 KI-KO* CD34^+^ HSPCs were sorted for double-positive tNGFR^+^/GFP^+^ expression (HDR values in Figure S4A) and sorted cells were subjected to IVTD. To validate *RAG2* biallelic editing efficiency, targeted alleles were analyzed by ddPCR and found to be positive (Figure S4B). Additionally, quantitative real-time PCR (qRT-PCR) analysis revealed that the endogenous gene expression of *RAG2* was markedly reduced (Figure S4C) while robust diverged cDNA expression was found exclusively in the engineered cells (Figure S4D). Importantly, the expression of the diverged *RAG2* cDNA indeed facilitated T-cell development highlighted by the successful differentiation of *RAG2 KI-KO* edited cells into CD3^+^ T cells relative to *RAG2* KO cells on day 28 (*RAG2 KI-KO*: 2.7%; *RAG2* KO: 0.03%) (Figure 3B-C, Figure S4E-F).

Deep-sequencing analysis of TRG V(D)J recombination on days 14 and 28 revealed a diverse TRG V(D)J rearrangement repertoire in the *RAG2 KI-KO* population following robust expression of the *RAG2* diverged cDNA (Figure 4A). Moreover, calculation of Shannon and Simpson diversity indices revealed that there were no significant differences between the TRG clonotype diversity richness of *Mock*, *CCR5* KO, and *RAG2 KI-KO* populations (Figure 4B). Lastly, the frequency distribution of complementarity determining region 3 (CDR3) lengths was comparable to that of the control groups, indicating production of a diverse TCR repertoire (Figure S5). The CDR3 region is responsible for recognizing processed antigen peptides and the length and sequence of the CDR3 varies by T-cell clone. Thus, the sequence of CDR3 determines the structure and specificity of the TCR, where a unique CDR3 sequence represents a specific T-cell clonotype. Sequencing of the CDR3 region can, therefore, be used as a measurement of TCR diversity^53^. Together, these data indicate that *KI* of the correction diverged cDNA promotes differentiation into CD3^+^ T cells and promotes the development of a highly diverse TRG repertoire.

**Figure 4.**
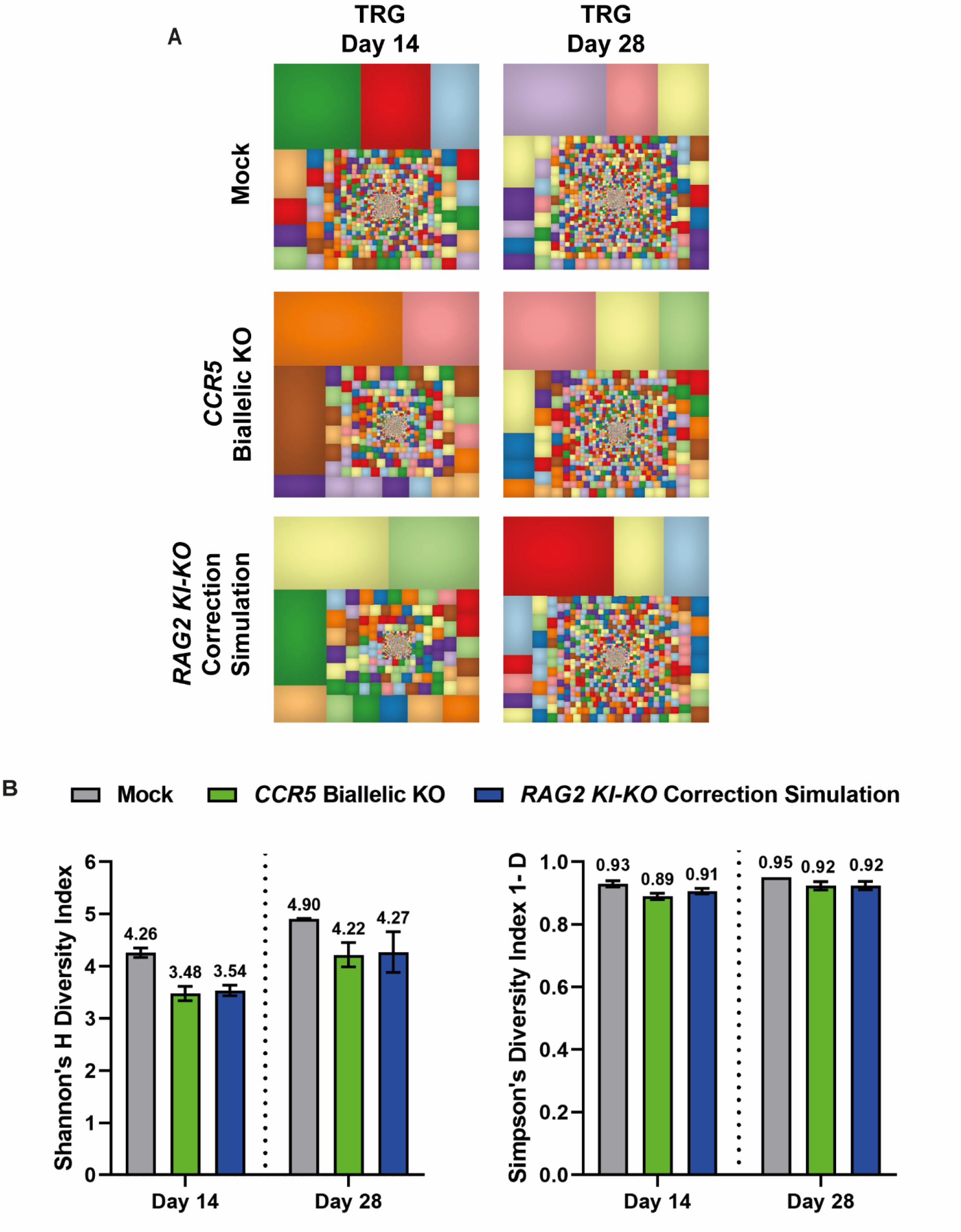
Expression of diverged *RAG2* cDNA in *KI-KO* cells leads to development of normal TCR repertoire. **(A)** Representative tree map depiction of the clonal complexity of the TRG deep-sequencing repertoire of differentiated T cells from *Mock*, *CCR5* biallelic KO, and *RAG2 KI-KO* correction simulation groups (due to lack of CD3 expression in the *RAG* biallelic KO population, sequencing and mapping of the repertoire was impossible). Each square represents a unique V-J pair and the size of each square represents the clone’s frequency. **(B)** Shannon and Simpson diversity indices of TRG repertoire on days 14 and 28 of *Mock*, *CCR5* biallelic KO, and *RAG2 KI-KO* correction simulation groups. (N=3). Data are represented as mean ± SEM.

### *RAG2* functional gene correction in *RAG2*-SCID-patient-derived CD34^+^ HSPCs

After determining that integration and expression of our correction donor was effective in facilitating T-cell development, we obtained CD34^+^ HSPCs from cord blood (CB) of a *RAG2*-SCID patient to actuate *in vitro* functional gene correction. This patient presented with a clinical SCID phenotype (Omenn phenotype) and had compound heterozygous missense mutations (G95V+E480X) in the *RAG2* gene^54^. In compound heterozygotes, each allele of the gene has a different genetic mutation. In such cases, our approach is uniquely advantageous since we integrate the complete *RAG2* cDNA including the endogenous 3’ UTR region at the initiation codon of the endogenous *RAG2* locus, thus resolving all possible mutations in the open reading frame while preserving the endogenous regulation and gene expression with a single rAAV6 donor.

Consequently, we targeted the *RAG2*-SCID CD34^+^ HSPCs with *RAG2* diverged cDNA (schematic depicted in Figure 3A) using a rAAV6 MOI of 12,500 viral genomes (VG) per cell (Figure S6A-B). An additional sample was exposed to a MOI of 6,250 VG/cell (Figure S6B) in an effort to minimize the potential for rAAV6-induced toxicity based on our group’s previous work that showed that rAAV6 vectors are recognized by the cells’ repair proteins leading to a DDR proportional to that of the MOI used^42^. Subsequently, cell sorting was used to enrich for *RAG2*-SCID corrected CD34^+^ HSPCs using the tNGFR marker and we subjected the sorted cells to the IVTD system. In parallel, HD-derived *Mock* CD34^+^ HSPCs, untreated *RAG2*-SCID CD34^+^ HSPCs, and unsorted *RAG2*-SCID corrected CD34^+^ HSPCs (MOI of 12,500 VG/cell) were cultured in the IVTD system. ddPCR was performed to determine the extent of *RAG2* cDNA genomic integration which was observed to be ~50% of all targeted alleles for both MOIs (Figure S6C). Following 28 days in the IVTD system, the *RAG2*-SCID corrected cells developed into CD3^+^ cells, in all three corrective populations, compared to *RAG2*-SCID cells that showed impaired progression towards CD3 expression (12,500 *RAG2*-SCID unsorted: 2.7%; 12,500 *RAG2*-SCID sorted: 6.0%; 6,250 *RAG2*-SCID sorted: 6.2%; *RAG2*-SCID: 0.2%) (Figure 5). These data were comparable with CD3-expression levels of *Mock* cells (average of 5%) after 28 days in the IVTD system (Figure 5 and S6D). The percentage of CD3^+^ cells in the population continued to rise by day 42 in the sorted populations similar to the *Mock* sample (Figure 5 and S6D). Additionally, cell viability staining showed an increase in cell death in the rAAV-treated populations by day 42, a hallmark of rAAV-induced toxicity (Figure S6E).

**Figure 5.**
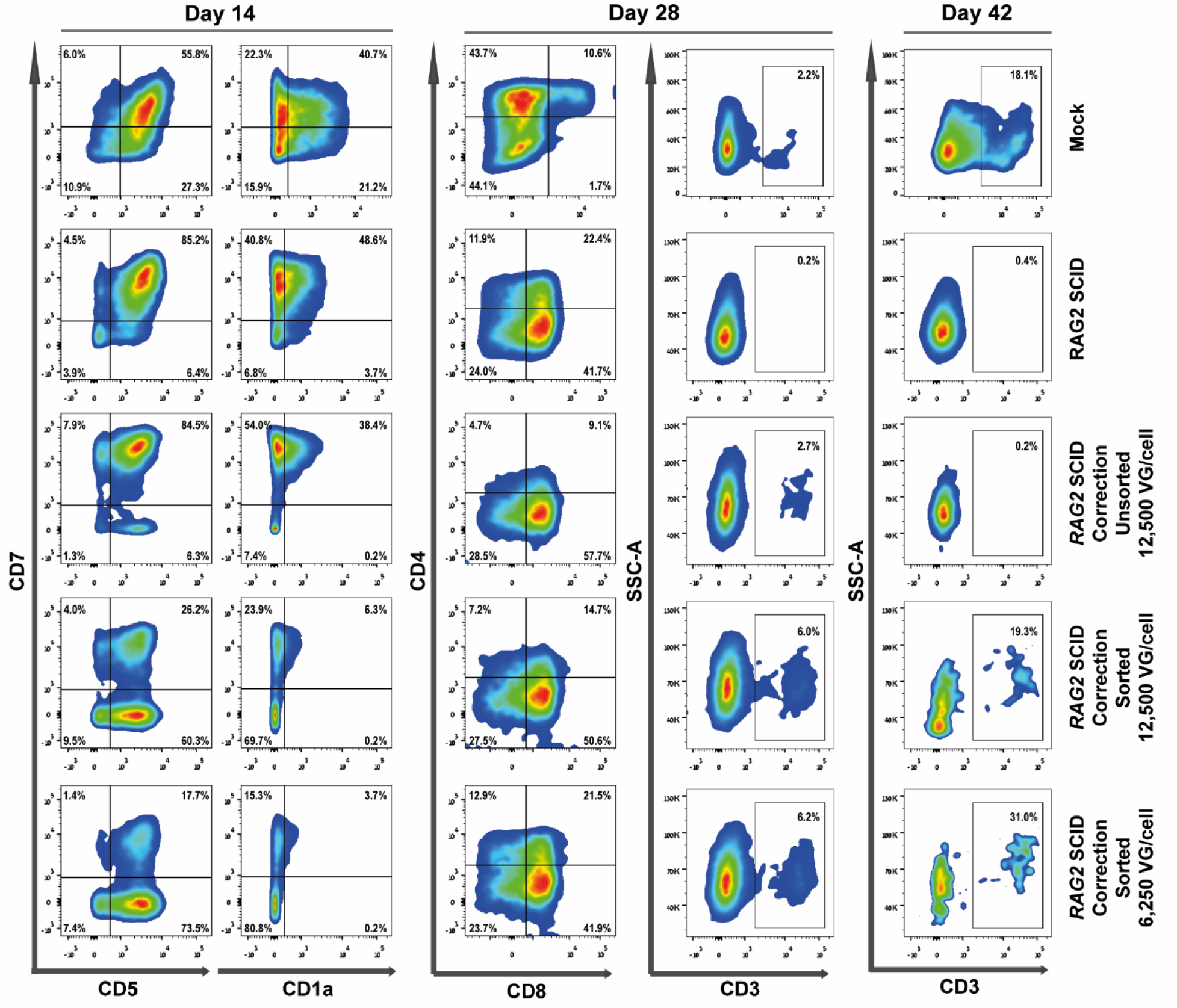
*RAG2* gene correction in *RAG2*-SCID patient-derived CD34^+^ HSPCs facilitates normal T-cell differentiation. Flow cytometry analysis of T-cell developmental progression, on days 14, 28, and 42, of the following cell groups: *Mock*, *RAG2*-SCID, unsorted *RAG2*-SCID correction with 12,500 VG/cell, sorted *RAG2*-SCID correction with 12,500 VG/cell, and sorted *RAG2*-SCID correction with 6,250 VG/cell. Corrected cells were treated with the same rAAV6 *RAG2* correction donor depicted in *Figure 3A*. Robust CD3 expression was found on day 28 in all *RAG2* correction populations, and on day 42 only in the sorted populations. Due to the rarity of patient-derived sample, N=1 for the *RAG2*-SCID samples and the *Mock* plots are from a representative donor.

Deep sequencing of the TRG repertoire revealed that each of the two sorted populations developed a diverse TRG repertoire, highlighted throughout the IVTD timeline by treemap, CDR3 length, and Shannon and Simpson diversity indices (Figure 6A-B and Figure S7). We could not, however, evaluate the repertoire of the unsorted *RAG2*-SCID correction cells at day 42 since only a very small number of CD3^+^ cells survived until this time point (Figure 5). The richness of the observed TRG repertoires were comparable to that of the untreated *Mock* populations in the IVTD system (Figure 6B). To further examine the rearrangement capability, copies of T-cell receptor excision circles (TRECs) were quantified. TRECs, are a DNA marker that represents the excision of the delta-coding segments out of the T-cell receptor alpha (TRA) locus, allowing the TRA recombination to occur^55^. According to newborn screening data for SCID, TREC copies are completely undetectable in *RAG2*-SCID samples^4^. In our samples, TRECs were detected only in the sorted *RAG2*-SCID correction samples at day 28 and 42 of differentiation, with a higher copy number in the 6,250 *RAG2*-SCID corrected cells (Table 1). In summary, building off of our success with our correction donor in the *KI-KO* correction simulation approach, we were able to utilize the same donor to correct *RAG2*-SCID-patient-derived CD34^+^ HSPCs *in vitro*, producing CD3^+^ T cells with diverse TRG repertoires.

**Table 1.** Quantitative PCR analysis of TRECs in *RAG2*-SCID, and *RAG2*-SCID correction cells through the over the 42 days in the IVTD system (days 14, 28, and 42).

|  |  | TREC<br>(copies) | RNase P<br>(average C<sub>T</sub> values) |
| --- | --- | --- | --- |
| Day 14 | *RAG2-SCID* | UN | 24.9 |
|  | Correction *RAG2-SCID* unsorted | UN | 24.8 |
|  | Correction *RAG2-SCID* sorted 12,500 VG/cell | UN | 24.3 |
|  | Correction *RAG2-SCID* with sorted 6,250 VG/cell | UN | 24.7 |
| Day 28 | *RAG2-SCID* | UN | 25.1 |
|  | Correction *RAG2-SCID* unsorted | UN | 24.7 |
|  | Correction *RAG2-SCID* sorted 12,500 VG/cell | 5.4 | 25.1 |
|  | Correction *RAG2-SCID* sorted 6,250 VG/cell | 22.4 | 24.9 |
| Day 42 | *RAG2-SCID* | UN | 24.7 |
|  | Correction *RAG2-SCID* unsorted | UN | 24.9 |
|  | Correction *RAG2-SCID* sorted 12,500 VG/cell | 8.3 | 25.6 |
|  | Correction *RAG2-SCID* sorted 6,250 VG/cell | 31.5 | 25.1 |

**Figure 6.**
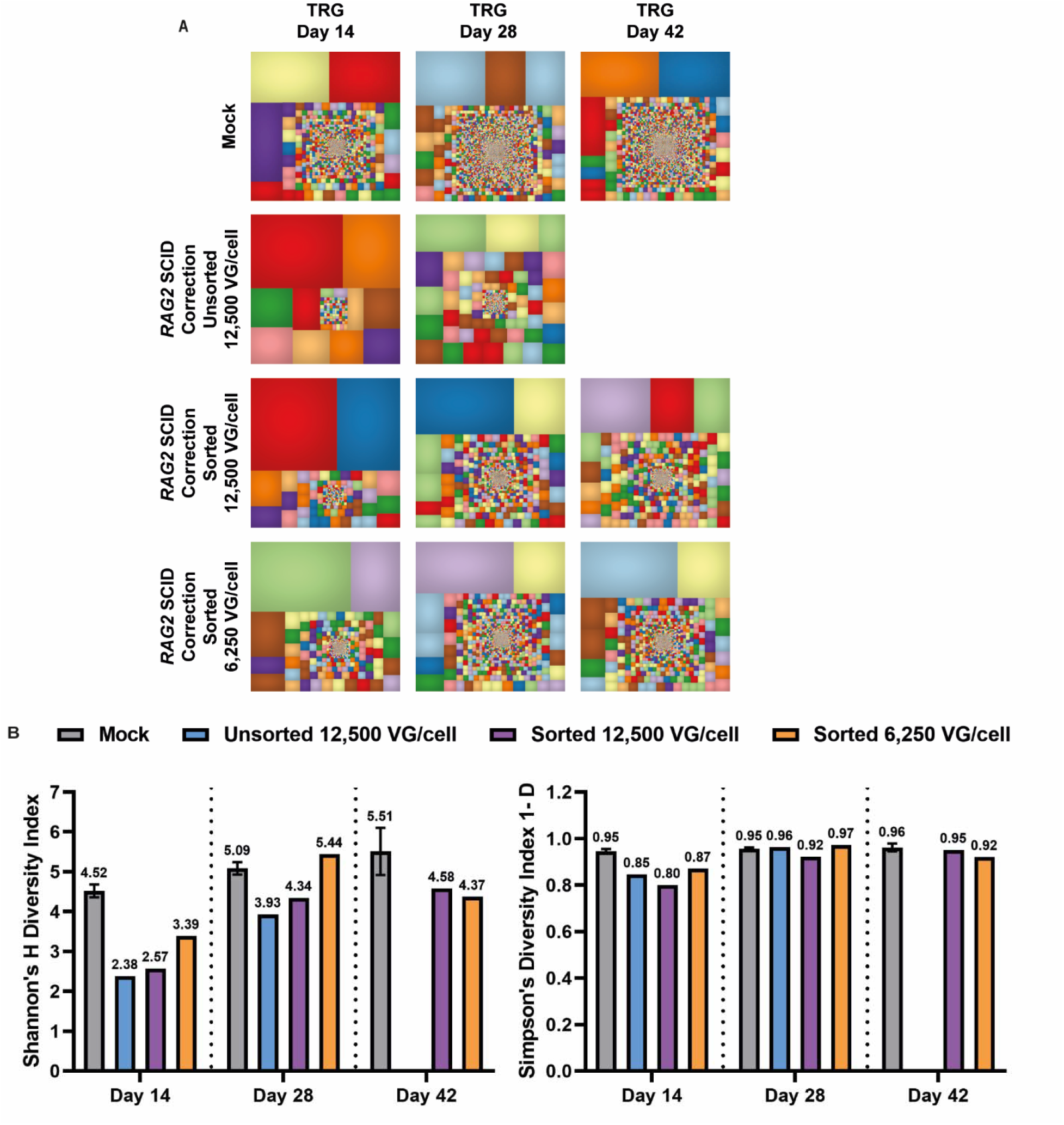
Corrected *RAG2* gene expression in *RAG2*-SCID patient-derived CD34^+^ HSPCs allows for normal TCR repertoire development. **(A)** Representative tree map depiction of the clonal complexity of the TRG deep-sequencing repertoire of differentiated T cells from *Mock* and *RAG2*-SCID correction populations, over the 42 days in the IVTD system (days 14, 28, and 42). **(B)** Shannon and Simpson diversity indices of TRG repertoires on days 14, 28, and 42 of *Mock* and *RAG2*-SCID correction groups (*Mock* [day 14 N=5, day 28 N=4, day 42 N=3]; and *RAG2*-SCID correction groups [day 14, day 28, and day 42 N=1]).

## Discussion

Disease modeling is a fundamental part of understanding the mechanisms underlying the expression and regulation of the affected genes and due to their central roles in the T-cell development process, we chose to focus on the *RAG1*, *RAG2*, *DCLRE1C*, and *IL7RA* genes. Patient-derived samples represent a natural system to study these mechanisms and their influence on the various cell processes, however, in diseases such as SCID, obtaining a sufficient quantity of CD34^+^ HSPCs from young patients for scientific studies is not a viable option. In place, some researchers have generated iPSCs derived from dermal fibroblasts or skin keratinocytes of SCID patients in order to build disease models and to study the cells’ differentiation potential^56–59^. This option has been widely accepted, however, iPSCs lack the authenticity of unadulterated CD34^+^ HSPCs. We present a novel approach to address this problem by modeling different recessive forms of SCID through biallelic CRISPR-Cas9/rAAV6-mediated gene *KO* in easily-attainable, abundant, biologically-authentic, HD-derived CD34^+^ HSPCs. This strategy is made possible by multiplexing HDR using distinct reporter genes and sorting via FACS to enrich for complete biallelic *KO* cells. While in this work biallelic HDR ranged from 0.7-6.3% (depending on the locus), we believe that enhancing HDR by reducing toxic rAAV6-induced DDR and/or inhibiting the NHEJ repair pathway, among other techniques, we can even further improve our method^60,61^.

We tracked the biallelic *KO* cells in a stromal cell-free IVTD system to better understand the SCID phenotype to subsequently use as confirmation for the success of our proof-of-concept correction phenotype. The readout of our system is multi-faceted, allowing for immunophenotyping via flow cytometry to track T-cell differentiation and NGS for TCR repertoire analysis. We believe that using this system for disease modeling via CRISPR-Cas9 editing has the potential to rapidly accelerate research on suspected disease-causing genes. These disease models can then be phenotypically and functionally validated in various differentiation platforms paving the way for new potential gene therapies.

One of the challenges when modeling SCID disease is effectively inducing a reproducible T-cell differentiation phenotype^62^. A recent study by Pavel-Dinu *et al.* presented an approach for *IL2RG* gene correction on male HD-derived CD34^+^ HSPCs and SCID-X1 patient samples cultured in the *in vitro* OP9 T-cell differentiation system. A major drawback of the OP9 system is the difficulty to efficiently differentiate PB CD34^+^ HSPCs while obtaining a consistent differentiation pattern^63,62,64^. Hence, Pavel-Dinu *et al.* addressed this problem by assessing the differentiation pattern of *IL2RG* HDR-corrected CD34^+^ HSPCs after transplantation to NSG mice. Through enrichment of edited cells via sorting and subsequent use of a stromal cell-free IVTD system we have overcome the variability challenges encountered in other T-cell differentiation studies. Thus, we established a sustainable, versatile, and reproducible system for evaluating gene-correction efficiency and studying the mechanisms of recessive forms of SCID that allows for easy cell tracking, immunophenotyping, and V(D)J recombination analyses.

With the use of the IVTD system, we showed the successful differentiation of HD-derived CD34^+^ HSPCs to CD7^+^CD5^+^ cells and eventually to CD4, CD8, and CD3-expressing T cells, with diverse TRG repertoires. *RAG2* biallelic KO and *DCLRE1C* biallelic KO CD34^+^ HSPCs developed into CD7^+^CD5^+^ cells, however, did not develop further to express CD3 and did not exhibit TRG gene rearrangement. Previously, Bifsha *et al.* and Bosticardo *et al.* assessed the T-cell differentiation outcome of SCID-patient-derived PB CD34^+^ HSPCs via the OP9 IVTD system and the artificial thymic organoids (ATO) system, respectively^62,65^. In their work, the *RAG2* patient cells displayed a developmental failure at the same T-cell differentiation stage (CD3-expressing stage) as we observed in our disease models as well as in the SCID-patient-derived PB CD34^+^ HSPC samples. Curiously, clinical analysis of *RAG2*-SCID patients c.G104T and p.G95V+E480X showed expression of CD3 (Table S1 and S2) despite the lack of differentiation of the same patients’ CD34^+^ HSPCs into CD3^+^ cells in the IVTD system. Although patients with SCID typically present profoundly reduced total T cell counts, patients can sometimes display the presence of self-reactive cells and trans-placentally acquired maternal T cells^66^. This could explain the discrepancy between clinical blood analysis and the output of our IVTD method. Taken together, modeling recessive forms of SCID via CRISPR-Cas9/rAAV6 gene editing coupled with the use of the highly reproducible IVTD system^44^ can effectively limit the requirement for large quantities of scarce patient-derived samples.

As mentioned, we chose to focus on the *RAG1*, *RAG2, DCLRE1C,* and *IL7RA* genes which combined are responsible for 33% of all SCID cases^3^, while *DCLRE1C* and *IL7RA* mutations alone were responsible for 59.3% of all SCID cases in Israel^4,7^. Even when a HLA-matched donor is found and allogeneic HSCT is conducted, patient outcomes for *RAG*-SCIDs are significantly worse than for T-B+NK-SCIDs such as SCID-X1^67,68^. *RAG*-SCIDs present a T-/B-/NK+ phenotype and the presence of NK cells has been shown to mediate graft rejection in murine SCID models as well as in human observational studies^69^. Thus, development of other treatment options for these SCIDs such as autologous gene therapy is critical to eliminate the need for finding an HLA-matched donor and to limit the risk of graft rejection.

Although great strides have been made in the past few decades, gene therapy using viral vectors in its current form (γRV or LV) could be hazardous for highly controlled and regulated genes such as *RAG1* and *RAG2* due to the constitutive expression of the transgene. In these cases, this can result in genomic instability, and in some cases leukemia, as a result of expression outside of the precise developmental window^70–72^. A recent study by Miyazaki *et al.* demonstrated that the mechanism for developmental regulation of *RAG1/2* gene expression is via cis-elements surrounding the *RAG1/2* coding regions which control the spatial genomic organization inside the locus^73^. Additionally, Rommel *et al*. found that aberrantly functioning RAG1/2 promotes lymphocyte malignancy through the formation of translocations and/or deletions in cancer-causing genes^74^. To address this, we aimed to establish a gene-editing method that addresses gene regulation and integration specificity. In our study, the *RAG2* sgRNA used for gene correction was designed to induce a specific DSB several base pairs downstream to the gene’s ATG start codon and the diverged cDNA delivered by the rAAV6 included the gene’s endogenous 3’ UTR. This targeting strategy allows for the subsequent expression of the transgene to rely on the endogenous transcriptional and translational regulation. To ensure the safety of our gene-editing strategy the *RAG2* sgRNA’s off-target potential was previously thoroughly examined, ensuring that it edits only a small number of off-target sites with low editing frequency^75,76^. Our approach highlights the importance of safeguarding the regulatory elements that are located outside of the coding region and the importance of inducing precise integration of transgenic elements.

After establishing a novel disease model, we proceeded to evaluate a proof-of-concept CRISPR-Cas9/rAAV6 gene-correction strategy through our *KI-KO* approach in HD-derived CD34^+^ HSPCs. Since *RAG2*-SCID is a recessive disorder, correction of only one allele is required to have the patient develop a functional immune system, hence our use of a single-allelic *KI* while the other allele is *KO* (mimicking a mutated allele). Our correction simulation using *RAG2* diverged cDNA expression led to successful T-cell differentiation highlighted by successful TRG gene rearrangement and expression of T-cell markers, CD4, CD8, and CD3. This allowed us to test our correction donor on HD-derived CD34^+^ HSPCs before utilizing any precious patient-derived samples. With the confidence in our system, we then actuated the first-of-its-kind, successful, *in vitro* correction of *RAG2*-SCID-patient-derived CD34^+^ HSPCs, which, post-correction, produced CD3-expressing T cells with a diverse TRG repertoire.

While rAAV6 is commonly used to deliver the donor DNA in genome-editing experiments, our group has shown in Allen *et al.* that rAAV6 vectors can trigger a potentially toxic DDR after entering cells proportional to the MOI used^42^. In order for CRISPR/rAAV6 treatments to be implemented as a clinical therapy for the purpose of gene correction, reduction of the rAAV6 toxicity is required. We demonstrated that lowering the MOI (from 12,500 to 6,250 VG/cell) maintains therapeutically relevant HDR and allows for potential alleviation of rAAV6-induced genotoxicity as noted by higher levels of TREC copy numbers as well as higher CD3 expression in the 6,250 VG/cell sample (Figure 5 and Table 1). Additionally, we noted a difference between the sorted and unsorted 12,500 VG/cell populations on day 28 and day 42 in the frequency of CD3^+^ cells. We believe that this is due to lower input of corrected cells to our IVTD system as well as competition between the corrected and uncorrected cells in our IVTD system in the unsorted sample. Due to these challenges we observed a drop off in CD3 expression in the unsorted sample while the CD3-expressing population in the comparable sorted sample expanded from 6% on day 28 to 19% on day 42. Based on this new understanding of the effects of MOI and cell enrichment, we propose the following possible ways to counter this reduction in efficacy: 1) Improving HDR efficiency by inhibiting the NHEJ repair pathway via molecules such as i53 and/or DNA-PK inhibitors; 2) Transiently suppressing p53 to rescue the toxic DDR^60,61^; and/or 3) Enrichment of the corrected cell population via cell sorting. Additionally, we are currently working towards using good manufacturing practices (GMP) grade rAAV6 preparations, free of potential toxic impurities. This will enable us to expand our research to animal models and eventually to the clinic.

In summary, we present a disease model for SCID and proof-of-concept gene therapy using a combination of CRISPR-Cas9, rAAV6 donors, a reproducible cell-free IVTD system, and abundant HD-derived CD34^+^ HSPCs. This system allows for circumventing the difficulty of obtaining large amounts of patient samples while providing a valuable tool that will allow researchers to find the optimal gene-editing configurations (i.e. engineered nuclease and donor DNA) for SCID and other additional recessive blood disorders. Lastly, since we use authentic CD34^+^ HSPCs as opposed to iPSCs, cell lines, or cells that no longer maintain their stemness, we believe that the feasibility of translating our gene correction strategy to the clinic will be simpler highlighted by our novel, successful correction of *RAG2*-SCID-patient-derived CD34^+^ HSPCs.

## Materials and methods

### CD34^+^ HSPCs purification

CD34^+^ HSPCs were isolated from CB samples collected at Sheba Medical Center CB Bank under Institutional Review Board - approved protocols to obtain CB units for research purposes. Informed consent was signed indicating that cord blood specimens that are not suitable for banking will be used for research. Mononuclear cells (MNCs) were separated from fresh CB samples by Lymphoprep™ density gradient medium (STEMCELL Technologies Inc.). Frozen CB samples were thawed and treated with 5,000 U/ml of DNase I (Worthington). Using the CD34^+^ Microbead Kit Ultrapure (Miltenyi Biotec), CD34^+^ HSPCs were purified from the MNCs by CD34 labeling according to the manufacturer’s protocol. Enriched CD34^+^ HSPCs were stained with APC anti-human CD34 antibodies (clone 581, Biolegend, San Jose, CA, USA), and sample purity was assessed on an Accuri C6 flow cytometer (BD Biosciences, San Jose, CA, USA). Cells were cryopreserved in CryoStor^®^ CS10 medium (STEMCELL Technologies Inc.) or were cultured for 48 hours at a density of 2.5 × 10^5^ cells/ml in StemSpan SFEM II (STEMCELL Technologies Inc.) supplemented with Stem Cell Factor (SCF) (100◻ng/ml), Thrombopoietin (TPO) (100◻ng/ml), Fms-like tyrosine kinase 3 ligand (Flt3-Ligand) (100◻ng/ml), Interleukin 6 (IL-6) (100◻ng/ml), StemRegenin 1 (SR1) (0.75◻mM), UM171 (35◻nM) (STEMCELL Technologies Inc.), and 1% Penicillin/Streptomycin (Biological Industries Israel Beit Haemek LTD). Cells were cultured at 37°C, 5% CO_2_, and 5% O_2_. The PB samples were obtained at Sheba Medical Center under Institutional Review Board. Informed consent was signed to ensure that specimens will be used only for research purposes. CD34^+^ HSPCs from PB *RAG1* and *RAG2* SCID patients, and from healthy donors were isolated by negative selection using RossetaSep^®^ (STEMCELL Technologies Inc.). Cells from the fractionated plasma layer were enriched by Lymphoprep™ density gradient medium (STEMCELL Technologies Inc.) and subsequently stained and sorted for APC human CD34^+^ (clone 581, Biolegend) and FITC human CD45^low^ (clone HI30, BD Pharmingen) expression by Aria III cell sorter (BD Biosciences). Post-sort, cells were stained with PE/Cy7-CD7 (clone:CD7-6B7, BioLegend) and BV421-CD5 (clone: UCHT2, BioLegend) and were analyzed by LSRFortessa™ cell analyzer (BD Biosciences).

### rAAV6 donor DNA template design

All rAAV6 vector plasmids were cloned using NEBuilder^®^ HiFi DNA Assembly Master Mix (cat # E2621L, New England Biolabs (NEB) Inc.) into the pAAV-MCS plasmid (Agilent Technologies containing inverted terminal repeats (ITRs). Each rAAV6 disruption donor contains a different reporter gene (GFP, tNGFR, or tdTomato) under the control of a SFFV promoter and followed by a BGH polyA sequence. The *RAG2* correction rAAV6 donor was designed and contains diverged *RAG2* cDNA followed by the *RAG2* 3’UTR and a reporter gene (tNGFR) controlled by a PGK promotor and BGH polyA sequence. Each donor DNA was designed with 400bp left and right homology arms flanking the sgRNA cut site. The rAAV6 vectors were produced by Vigene Biosciences in large-scale rAAV6 packaging. See Table S5 for the DNA donor sequences.

### CRISPR-Cas9 genome targeting and biallelic integration

*RAG2* and *CCR5* modified sgRNAs, previously described^37,75^, were synthesized by TriLink BioTechnologies. *IL7RA* and *DCLRE1C* modified sgRNAs were synthesized by Integrated DNA Technologies (IDT, Coralville, IA). The 20bp sgRNA genomic target sequences were as follows: *RAG2*: 5’-UGAGAAGCCUGGCUGAAUUA-3’, *CCR5*: 5’-GCAGCAUAGUGAGCCCAGAAG-3’, *IL7RA*: 5’-ACAAUUCUAGGUACAACUUU-3’, and *DCLRE1C*: 5’-GCGCUAUGAGUUCUUUCGAG-3’. 260 pmol of sgRNA was complexed pre-electroporation with 104 pmol of Alt-R Cas9 protein (IDT, Coralville, IA) forming an RNP complex, at a 1:2.5 molar ratio (Cas9:sgRNA). Electroporation of CD34^+^ HSPCs was performed with P3 nucleofection solution (Lonza, Basel, Switzerland) in the Lonza Nucleofector 4D (program DZ-100). Electroporated CD34^+^ HSPCs were plated at 4 × 10^5^ cells/ml and transduction of disruption *RAG2* donors was performed at a multiplicity of infection (MOI) of 6,250 VG/cell for each donor. The MOI for the *RAG2* correction rAAV6 in HD-derived CD34^+^ HSPCs was 12,500 VG/cell, and for *RAG2*-SCID CD34^+^ HSPCs the MOI was 6,250 or 12,500 VG/cell. For *IL7RA* rAAV6 disruption donors, GFP and tdTomato, MOIs were 12,500 VG/cell and 25,000 VG/cell, respectively. For *DCLRE1C* rAAV6 disruption donors, GFP and tdTomato, the MOI was 12,500 VG/cell for each donor. rAAV6 donors were added to the plated cells, within 15 minutes of electroporation. After 24 hours, fresh CD34^+^ medium was added to form a final concentration of 2.5 × 10^5^ cells/ml. 48 hours post-electroporation, cells were collected and prepared for the biallelic enrichment by the Aria III cell sorter (BD Biosciences). Cells were stained with PE human CD34 (clone: 561, BioLegend) antibodies, and when rAAV6 tNGFR is transduced APC-tNGFR (clone: ME20.4, BioLegend) staining is performed as well.

### Digital Droplet PCR™

The quantification of genomic integration was performed by Digital Droplet PCR™ (ddPCR™, Bio-Rad, Hercules, CA, USA). Genomic DNA was extracted from sorted populations cultured in CD34^+^ medium and IVTD using GeneJET Genomic DNA Purification Kit (Thermo Fisher Scientific, USA). Each ddPCR reaction mix contains a HEX reference assay detecting copy number input of the *CCRL2* gene to quantify the chromosome 3 input. Separate assays with a set of target-specific primers and a FAM-labeled probe (500◻nM and 250◻nM, respectively) for each gene were used to detect the locus specific donor integration. The reaction was carried as follows: 2X ddPCR Supermix for Probes No dUTP (Bio-Rad), 1X of each PrimeTime^®^ Standard qPCR Assay (IDT, Coralville, IA), 1X of a restriction enzyme mix [1/2 EcoRI-HF^®^ (NEB #R3101, 2/5 Nuclease-free water, 1/10 CutSmart Buffer 10X (NEB)], 20 ng genomic template DNA, and supplemented to a total of 20 μl with Nuclease-free water. Droplet samples were prepared according to manufacturer’s protocol (Bio-Rad) and 40◻μl of the droplets conveyed to a 96-well plate and amplified in a Bio-Rad PCR thermocycler with the following PCR conditions: 1 cycle at 95°C for 10 minutes, followed by 40 cycles at 95°C for 30 seconds and then 55°C for 3 minutes, followed by 1 cycle at 98°C for 10 minutes at a ramp rate of 2.2°C/s. After the PCR, the 96-well plate was loaded in the QX200 droplet reader (Bio-Rad). The droplets from each well were analyzed and the concentration of copies/μl of the site-specific donor integration (FAM) and wild-type *CCRL2* (HEX) alleles were calculated using the QuantaSoft analysis software (Bio-Rad). Primers and probes sequences are presented in Table S3.

### Estimation of mRNA levels in differentiated T cells

RNA was extracted using Direct-zol™ RNA Miniprep Plus (Zymo Research, cat #R2073) from differentiated T cells obtained on days 14 and 28 of IVTD. cDNA was prepared from 50-150 ng RNA, using Oligo d(T)23 VN-S1327S (NEB), dNTPs 10mM, and M-MuLV Reverse Transcriptase (cat# M0253S, NEB). qRT-PCR reactions were done using TaqMan® Fast Advanced Master Mix (cat# AB-4444557, Thermo Fisher Scientific) and carried out on StepOnePlus™ Real-Time PCR System (cat# 4376600, Thermo Fisher Scientific). PCR conditions were as follows: uracil-N-glycosylase gene (UNG) incubation was 2 minutes at 50°C, polymerase activation was 20 seconds at 95°C, followed by 40 cycles of 1 second at 95°C and 20 seconds at 60°C. Primers and probes sequences are presented in Table S4.

### *In vitro* T-cell differentiation (IVTD) system

Using StemSpan™ T Cell Generation Kit (STEMCELL Technologies, Inc.), CD34^+^ HSPCs were cultured in StemSpan™ SFEM II containing Lymphoid Progenitor Expansion Supplement on plates coated with Lymphoid Differentiation Coating Material for 14 days. Subsequently, cells were harvested and re-seeded on coated plates with StemSpan™ T Cell Progenitor Maturation Supplement for an additional 14 days. PB and CB CD34^+^ HSPCs experiments, were re-seeded on coated plates with StemSpan™ T Cell Progenitor Maturation Supplement at day 28 of culturing and harvested at day 42 of differentiation. Flow cytometry analysis was conducted at each time point of differentiation using the LSRFortessa™ cell analyzer (BD Biosciences). On day 14, collected cells were stained with PE/Cy7-CD7 (clone:CD7-6B7, BioLegend), BV421-CD5 (clone: UCHT2, BioLegend), and CD1a-PE (clone: BL6, Beckman Coulter, USA) or CD1a-APC (clone: BL6, Beckman Coulter, USA) antibodies. On days 28 and 42, collected cells were stained with PE/Cy7-CD4 (clone: RPA-T4, BioLegend), APC-r700-CD8a (clone: RPA-T8, BD Horizon™), and BV421-CD3 (clone: UCHT1, BioLegend) antibodies. BD Horizon™ Fixable Viability Stain 510 was performed on all the collected cells at each time point. APC-NGFR (clone: ME20.4, BioLegend) staining was conducted on all tNGFR rAAV6 integrated cells. For evaluating the background staining, fluorescence minus one (FMO) + isotype control antibody staining was performed using the following isotypes: PE/Cy7 Mouse IgG2a κ (cat # 400232, BioLegend), BV-421 Mouse IgG1 κ (cat # 400158, BioLegend), PE Mouse IgG1 κ (cat # 400112, BioLegend), PE/Cy7 Mouse IgG1 κ (cat # 400126, BioLegend), APC-R700 Mouse IgG1 κ (cat #564974, BD Horizon™), and APC Mouse IgG1 k (cat # 400122, BioLegend).

### Identification of TRG gene rearrangements

gDNA from differentiated T cells and CD34^+^ HSPCs was extracted by the GeneJET Genomic DNA Purification Kit (Thermo Fisher Scientific). TRG rearrangement was assessed by PCR amplification of 12 possible CDR3 clones, using combinations of 4 primers for Vγ and 3 primers for Jγ regions in each reaction (IdentiClone™ TCRG Gene Clonality Assay, Invivoscribe, Inc.). TRG clonality was ran and analyzed on 2% agarose gel. For deep sequencing of the TRG repertoire, the TRG rearranged genomic products were amplified using a single multiplex master mix LymphoTrack^®^ TRG assay (Invivoscribe, Inc.). PCR amplicons were purified and sequenced using the Miseq V2 (500 cycles) kit, 250-bp paired-end reads (Illumina, San Diego, CA). FASTQ files were analyzed by LymphoTrack Software (Invivoscribe, Inc.). LymphoTrack Software unique sequences generated data of each sample furthered analyzed by IMGT^®^, the international ImMunoGeneTics information system^®^ (HighV-QUEST, http://www.imgt.org). The analysis of the incidence of productive and unproductive TRG rearrangements sequences were performed amidst the total sequences and presented visually by TreeMap 2019.3.2 software (Macrofocus GmbH). Unique sequences and CDR3 length analysis were obtained from the total productive sequences. The Shannon’s H and Simpson’s D diversity indices were calculated by PAST software^77^. The sequencing data were deposited to the Sequence Read Archive (SRA), under accession number: PRJNA838341.

### Quantification of TRECs

TREC copy numbers were analyzed by qPCR, as previously described^78^. DNA samples were examined in triplicates, 100 ng for each replicate. qPCR reactions were carried out on StepOnePlus™ Sequence Detector System (Applied Biosystems). A standard curve was assembled by utilizing serial dilutions with 10^3^-10^6^ copies of TREC plasmid. RNase P amplification (TaqMan assay, Applied Biosystems) was conducted for quality control of the DNA TREC amplification.

## Supporting information

Hendel Supplemental data

## Acknowledgments

We would like to thank the members of the Somech and the Hendel Labs for reading the manuscript and providing practical advice. Additionally, we want to thank Dr. Rasmus O. Bak, and Dr. Andreas Reinisch for numerous important discussions and comments. Lastly, we would like to thank D. Russell for providing the pDGM6 plasmid. This work was supported by the European Research Council (ERC) under the European Union Horizon 2020 research and innovation program (grant number. 755758, A.H.), Israel Science Foundation (grant number. 2031/19, A.H.), and Israel Precision Medicine Program (grant number. 3115/19, R.S., A.H., and Y.N.L). Figure S2A was created with BioRender.com.

## Author Contributions

O.I., D.A., O.K., Y.Z., D.B., A.A., and A.L. designed and conducted the experiments, evaluated, and analyzed the data; O.I. and D.B. performed the bioinformatics analyses, with the help and guidance of Y.N.L.; K.B., and A.N. provided cord blood samples and R.S. provided PB samples; K.B., Y.N.L., A.N., and R.S. critically reviewed the experiments and provided important advice; A.H. supervised and conceived the research and planned the experiments and the approaches; A.H., D.A., O.I., and Y.Z. wrote the manuscript, with contributions from all authors.

## Supplemental information

Supplemental data. Figures S1–S7; Tables S1-S4.

Table S5. rAAV6 donor DNA sequences.

