## Supplementary material for "Multiplex HDR for Disease and Correction Modeling of SCID by CRISPR Genome Editing in Human HSPCs": Hendel Supplemental data

### Supplementary figures-

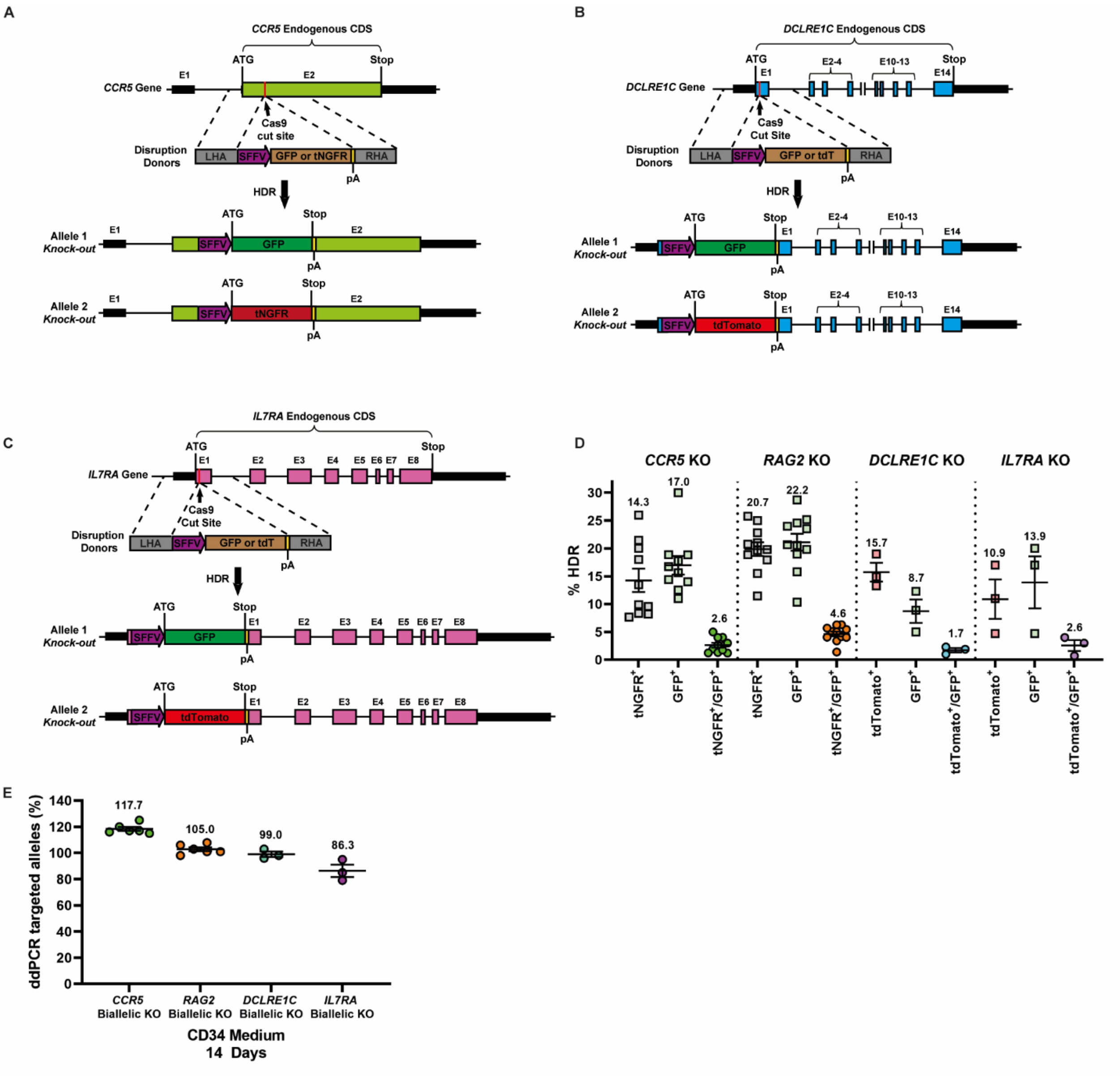


**Figure S1. Biallelic targeting of SCID-related genes via CRISPR-Cas9/rAAV6 in HD-derived CD34$^+$ HSPCs. (A, B, and C)** Schematic representations of **(A)** *CCR5*, **(B)** *DCLRE1C*, and **(C)** *IL7RA* disruption donors containing a tNGFR, GFP, or tdTomato selectable marker driven by a SFFV promoter, and two arms of homology, centered at the CRISPR-Cas9 cut site, for direct recombination repair (left- LHA, right- RHA). (**D**) HDR efficiencies as measured by flow cytometry for *CCR5* KO, *RAG2* KO, *DCLRE1C* KO, and *IL7RA* KO (N=10, N=11, N=3, and N=3, respectively). The HDR efficiencies of the three disruption donors alone (tNGFR$^+$, GFP$^+$, or tdTomato$^+$) as well as the multiplex HDR efficiencies (tNGFR$^+$/GFP$^+$ or tdTomato$^+$/GFP$^+$) are presented for each locus. **(E)** Site-specific HDR efficiencies at the *CCR5*, *RAG2*, *DCLRE1C*, and *IL7RA* loci measured by ddPCR, and normalized by targeted *CCRL2* alleles. Genomic DNA was extracted from undifferentiated CD34$^+$ HSPCs that were sorted for biallelic *KO* donor integration. (*CCR5* and *RAG2* [N=6] and *DCLRE1C* and *IL7RA* [N=3]). Data are represented as mean ± SEM.

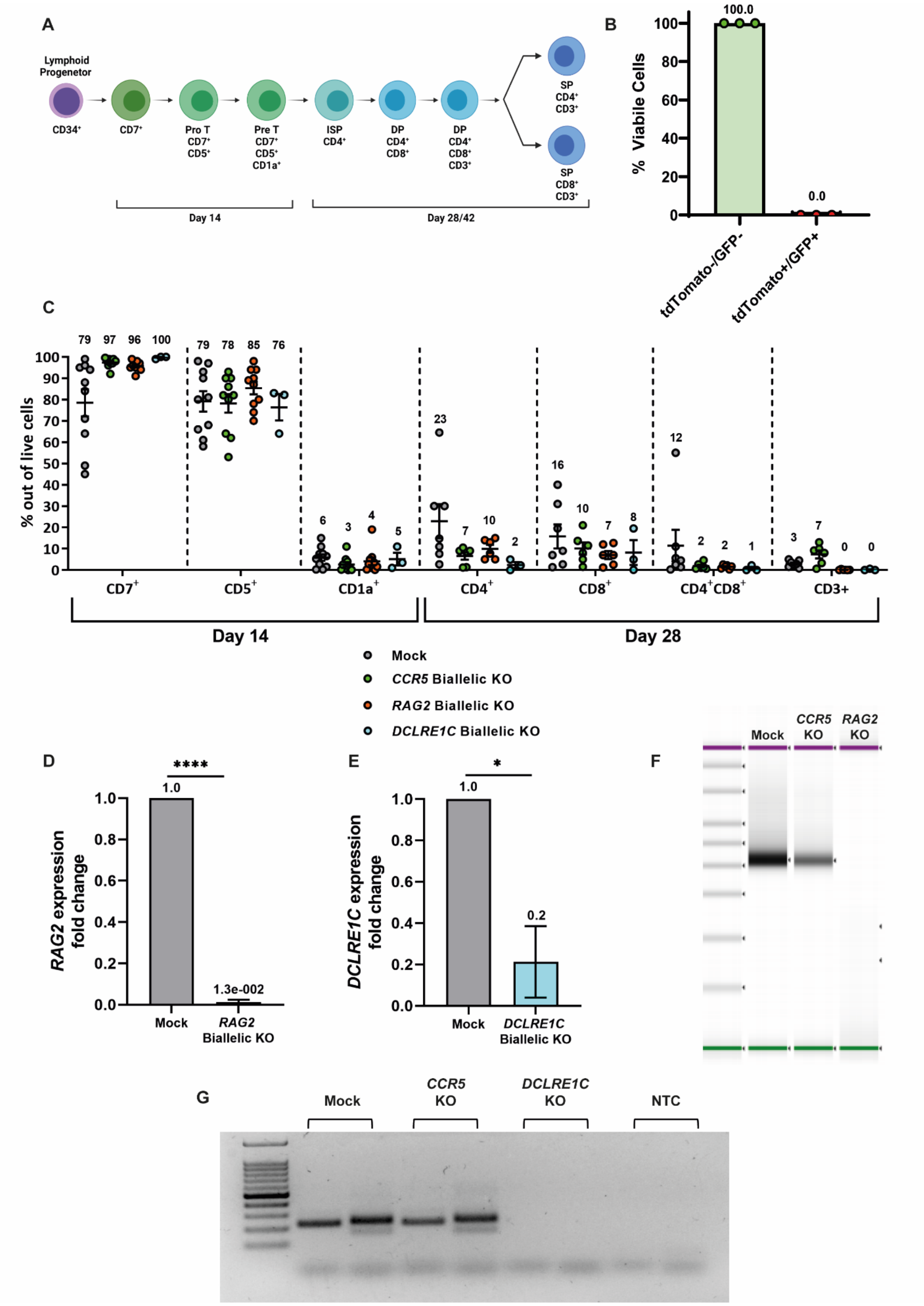
A
Lymphoid Progenitor
CD34+
CD7+
Pro T CD7+ CD5+
Pre T CD7+ CD5+ CD1a+
ISP CD4+
DP CD4+ CD8+
DP CD4+ CD8+ CD3+
SP CD4+ CD3+
SP CD8+ CD3+
Day 14
Day 28/42
B
% Viabile Cells
100.0
0.0
tdTomato-/GFP-
tdTomato+/GFP+
C
% out of live cells
CD7+
CD5+
CD1a+
CD4+
CD8+
CD4+CD8+
CD3+
Day 14
Day 28
Mock
CCR5 Biallelic KO
RAG2 Biallelic KO
DCLRE1C Biallelic KO
D
RAG2 expression fold change
****
1.0
1.3e-002
Mock
RAG2 Biallelic KO
E
DCLRE1C expression fold change
*
1.0
0.2
Mock
DCLRE1C Biallelic KO
F
Mock
CCR5 KO
RAG2 KO
G
Mock
CCR5 KO
DCLRE1C KO
NTC

15

**Figure S2. Modeling SCID abberant T-cell development phenotype via biallelic *KO* in HD-derived CD34$^+$ HSPCs in a cell-free IVTD assay. (A)** Differentiation of CD34$^+$ HSPCs to T cells in the IVTD system. HSPCs develop into CD7$^+$/CD5$^+$ pro-T cells which then continue on to develop into T-cell committed pre-T CD1a$^+$ cells. These CD7$^+$/CD5$^+$/CD1a$^+$ cells can then become CD4$^+$ ISP T cells. These ISPs give rise to DP CD4$^+$/CD8$^+$ cells that can begin to express CD3. These DP CD3$^+$ cells then undergo full maturation into SP CD8$^+$/CD3$^+$ or CD4$^+$/CD3$^+$ cells. **(B)** Distribution of viable cells based on genotype post-multiplex HDR. Cells were sorted for tdTomato$^+$/GFP$^+$ disruption donor integration at the *IL7RA* locus. After 6 days in the IVTD system viability staining was conducted and the double-negative (tdTomato$^-$/GFP$^-$) and double-positive (tdTomato$^+$/GFP$^+$) populations were analyzed independantly. **(C)** Summary of T-cell marker expression in *Mock*, *CCR5* biallelic KO, *RAG2* biallelic KO, and *DCLRE1C* KO populations on day 14 (N=10, N=10, N=10, and N=3, respectively) and on day 28 of IVTD (N=7, N=6, N=6, and N=3, respectively). For all *RAG2* biallelic KO and *DCLRE1C* biallelic KO samples, CD3 expression is <0.1. **(D)** Measurement of the endogenous *RAG2* gene expression in the *RAG2* biallelic KO and *Mock* cells at day 14 of IVTD (N=3) **(E)** Measurement of the endogenous *DCLRE1C* gene expression in the *DCLRE1C* biallelic KO and *Mock* cells at day 14 of IVTD (N=3). **(F)** TapeStation analysis of TRG repertoire libraries for deep sequencing of *Mock*, *CCR5* biallelic KO, and *RAG2* biallelic KO cells at day 14 of IVTD. **(G)** Examination of the TRG V(D)J recombination process by PCR analysis using primers that flank the V-J regions for the *Mock*, *CCR5* biallelic KO, and *DCLRE1C* biallelic KO cells at 14 days of IVTD. * $p<0.05$ and **** $p<.0001$ (*t*-test). Data are represented as mean ± SEM.

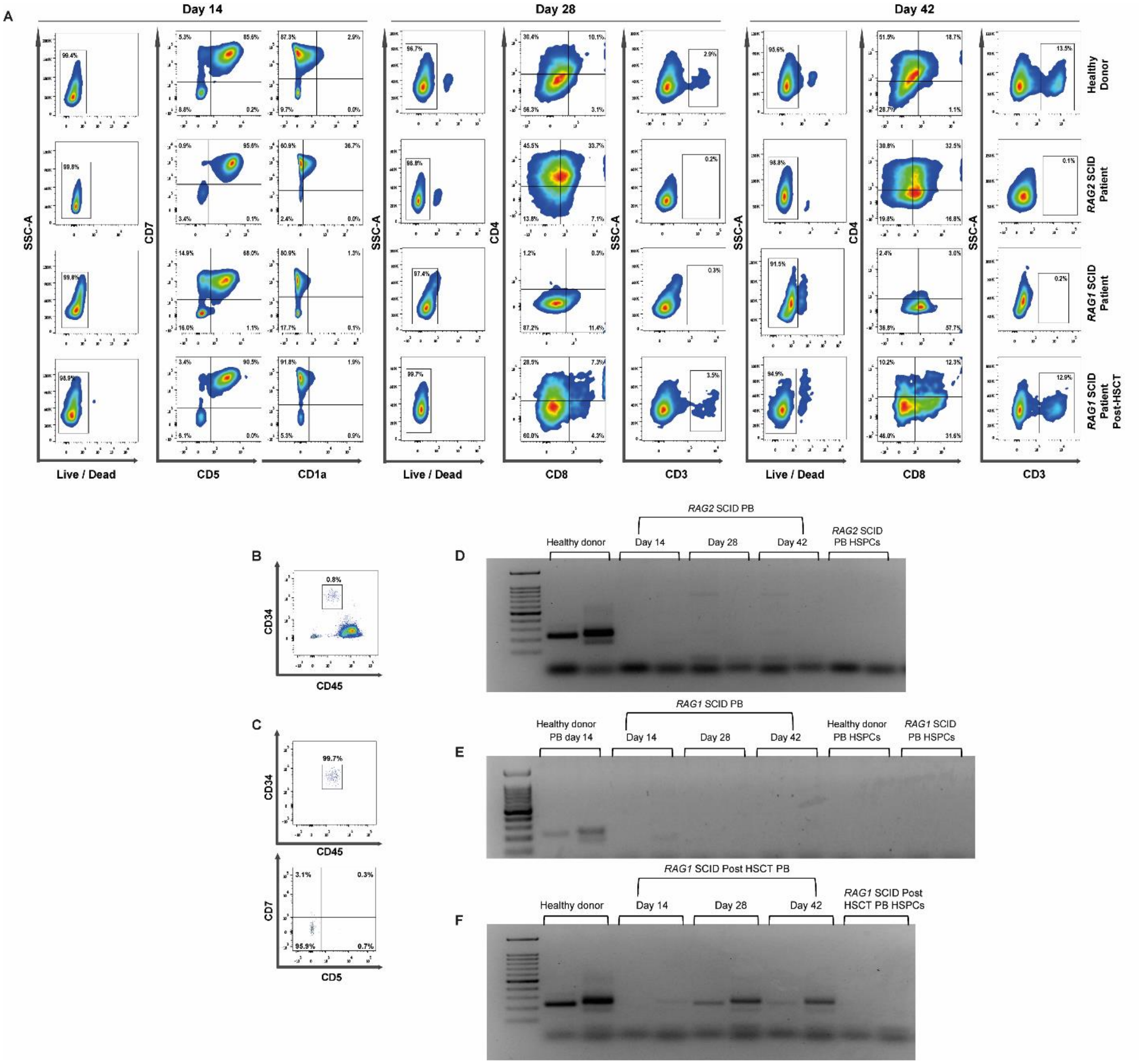


**Figure S3. SCID-patient-derived PB CD34⁺ HSPCs do not differentiate to CD3⁺ T Cells in the IVTD system. (A)** Flow cytometry analysis of the T-cell developmental progression of HD-derived, *RAG2*-SCID patient-derived, and *RAG1*-SCID-patient-derived PB CD34⁺ HSPCs, as well as *RAG1*-SCID-derived PB CD34⁺ HSPCs collected post HSCT samples. At day 14 of IVTD, CD7, CD5, and CD1a expression were examined. Following 28 and 42 days of IVTD, CD4, CD8, and CD3 T-cell marker expression were analyzed. Only the HD-derived and post HSCT *RAG1*-SCID patient-derived samples progressed to CD3⁺ stage of T-cell differentiation. Gating was determined by FMO + isotype controls. For viability staining of samples the gating determination is based on unstained cells. **(B)** Enrichment of PB CD34⁺ HSPC by sorting for human

CD34$^{high}$/CD45$^{low}$ expression. **(C)** Immediately after the sorting, CD34$^{+}$ HSPCs, were stained for CD7 and CD5 expression and re-analyzed for validation of CD34 and CD45 expression. The enriched cells were found to be T-cell free. **(D, E, and F)** Examination of the TRG V(D)J recombination of **(D)** *RAG2*-SCID patient-derived cells, **(E)** *RAG1*-SCID patient-derived cells, and **(F)** *RAG1*-SCID post HSCT patient-derived cells on days 14, 28, and 42 of the IVTD system. **(D)** The targeted samples are as follows (*left to right*): HD-derived PB CD34$^{+}$ HSPCs on day 14 of IVTD, *RAG2*-SCID-patient-derived PB CD34$^{+}$ HSPCs on days 14, 28, and 42 of the IVTD system, undifferentiated HD-derived CD34$^{+}$ HSPCs, and unedited and undifferentiated *RAG2*-SCID-patient-derived PB CD34$^{+}$ HSPCs. **(E)** The targeted samples are as follows (*left to right*): HD-derived PB CD34$^{+}$ HSPCs on day 14 of IVTD, *RAG1*-SCID-patient-derived PB CD34$^{+}$ HSPCs on days 14, 28, and 42 of the IVTD system, undifferentiated HD-derived CD34$^{+}$ HSPCs, and undifferentiated *RAG1*-SCID-patient-derived PB CD34$^{+}$ HSPCs. **(F)** The targeted samples are as follows (*left to right*): HD-derived PB CD34$^{+}$ HSPCs on day 14 of IVTD, *RAG1*-SCID post-HSCT patient-derived PB CD34$^{+}$ HSPCs on days 14, 28, and 42 of the IVTD system, and undifferentiated *RAG1*-SCID post-HSCT patient-derived PB CD34$^{+}$ HSPCs.

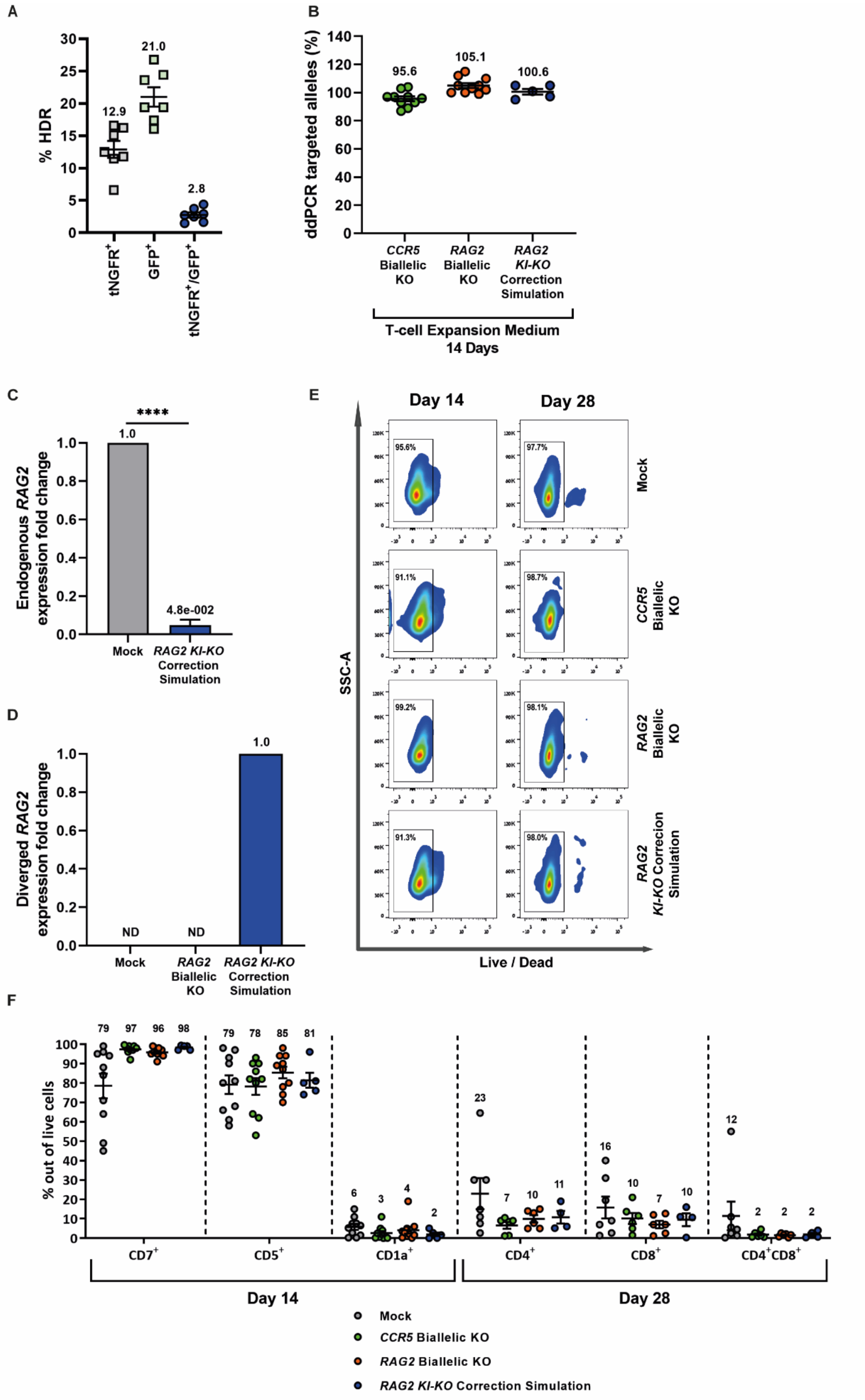
A
% HDR
12.9
21.0
2.8
tNGFR+
GFP+
tNGFR+/GFP+
B
ddPCR targeted alleles (%)
95.6
105.1
100.6
CCR5 Biallelic KO
RAG2 Biallelic KO
RAG2 KI-KO Correction Simulation
T-cell Expansion Medium
14 Days
C
Endogenous RAG2 expression fold change
****
1.0
4.8e-002
Mock
RAG2 KI-KO Correction Simulation
D
Diverged RAG2 expression fold change
ND
ND
1.0
Mock
RAG2 Biallelic KO
RAG2 KI-KO Correction Simulation
E
Day 14
Day 28
95.6%
97.7%
Mock
91.1%
98.7%
CCR5 Biallelic KO
99.2%
98.1%
RAG2 Biallelic KO
91.3%
98.0%
RAG2 KI-KO Correcion Simulation
SSC-A
Live / Dead
F
% out of live cells
79
97
96
98
79
78
85
81
6
3
4
2
23
7
10
11
16
10
7
10
12
2
2
2
CD7+
CD5+
CD1a+
CD4+
CD8+
CD4+CD8+
Day 14
Day 28
Mock
CCR5 Biallelic KO
RAG2 Biallelic KO
RAG2 KI-KO Correction Simulation

62

**Figure S4. Proof-of-concept functional gene correction simulation via insertion of diverged *RAG2* cDNA in HD-derived CD34⁺ HSPCs. (A)** Multiplex HDR efficiencies for *RAG2 KI-KO* editing as measured by flow cytometry (N=7). **(B)** Site-specific HDR efficiencies at the *CCR5* or *RAG2* loci measured by ddPCR and normalized by targeted *CCRL2* alleles at day 14 of IVTD (*CCR5* and *RAG2* biallelic KO [N=10], and *RAG2 KI-KO* correction simulation [N=5]). **(C)** Measurement of endogenous *RAG2* gene expression in the *RAG2 KI-KO* correction simulation cells compared to *Mock* cells on day 14 of IVTD (N=3). **(D)** Measurement of *RAG2* diverged cDNA expression in the *Mock* and *RAG2* biallelic KO cells compared to *RAG2 KI-KO* correction simulation cells on day 14 of IVTD (N=3). ND = Not Detected. **(E)** Representative plots depicting viability staining of samples on days 14 and 28 of IVTD. Gating determination is based on unstained cells. **(F)** Summary of T-cell marker expression in *Mock*, *CCR5* biallelic KO, *RAG2* biallelic KO, and *RAG2 KI-KO* correction simulation populations on day 14 (N=10, N=10, N=10, and N=5, respectively) and on day 28 of IVTD (N=7, N=6, N=6, and N=4, respectively). The data for *Mock*, *CCR5* KO, and *RAG2* KO are taken from *Figure S2C* and are presented here for comparison to the *RAG2 KI-KO* correction simulation population. **** $p<0.0001$ (*t*-test). Data are represented as mean ± SEM.

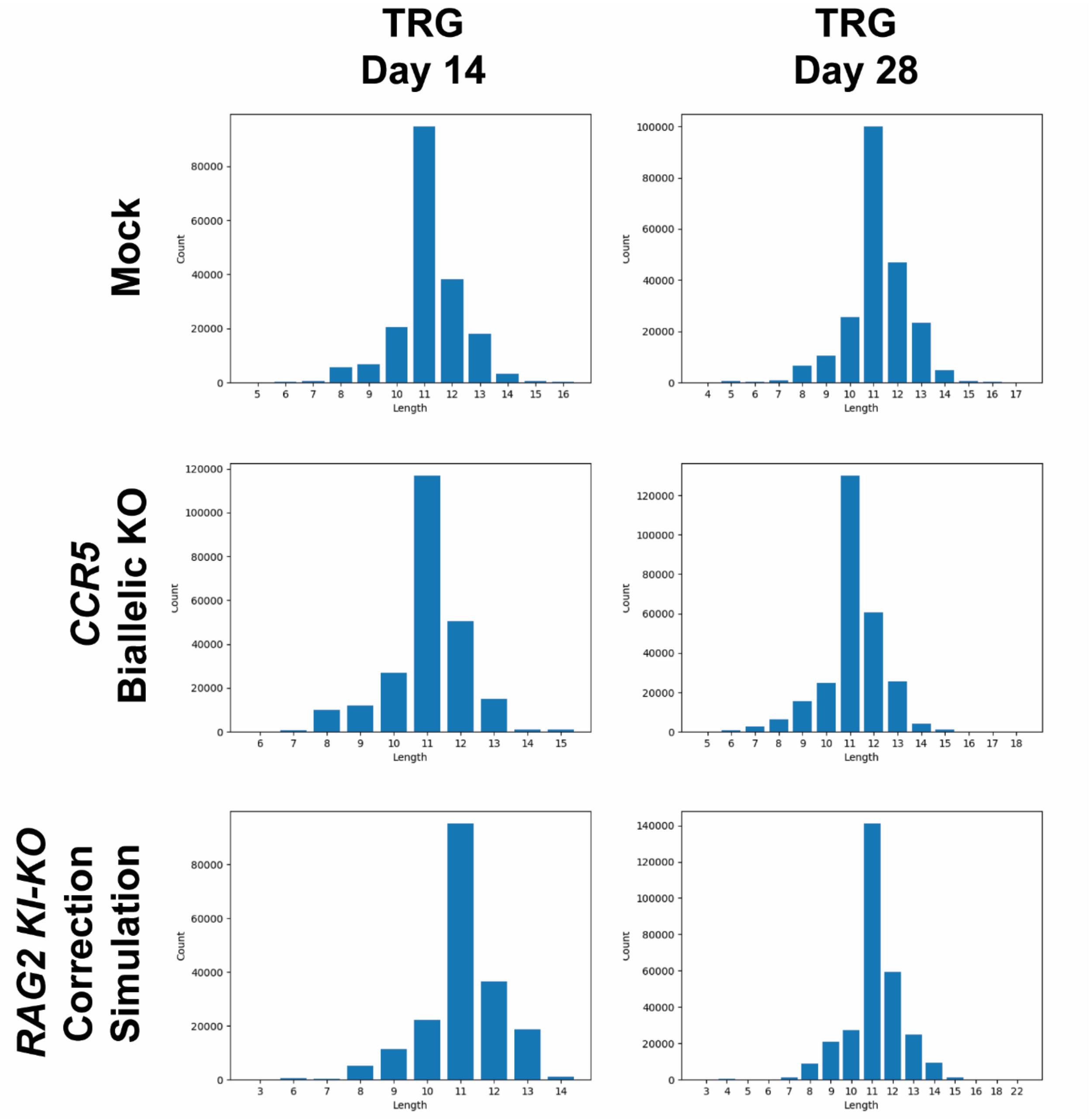


**Figure S5. Expression of *RAG2* diverged cDNA in *KI-KO* cells leads to normal TRG V(D)J recombination.** Representative CDR3 length distribution among unique TRG sequences expressed on days 14 and 28 in *CCR5* biallelic KO and *RAG2 KI-KO* correction simulation populations compared to their *Mock* group.

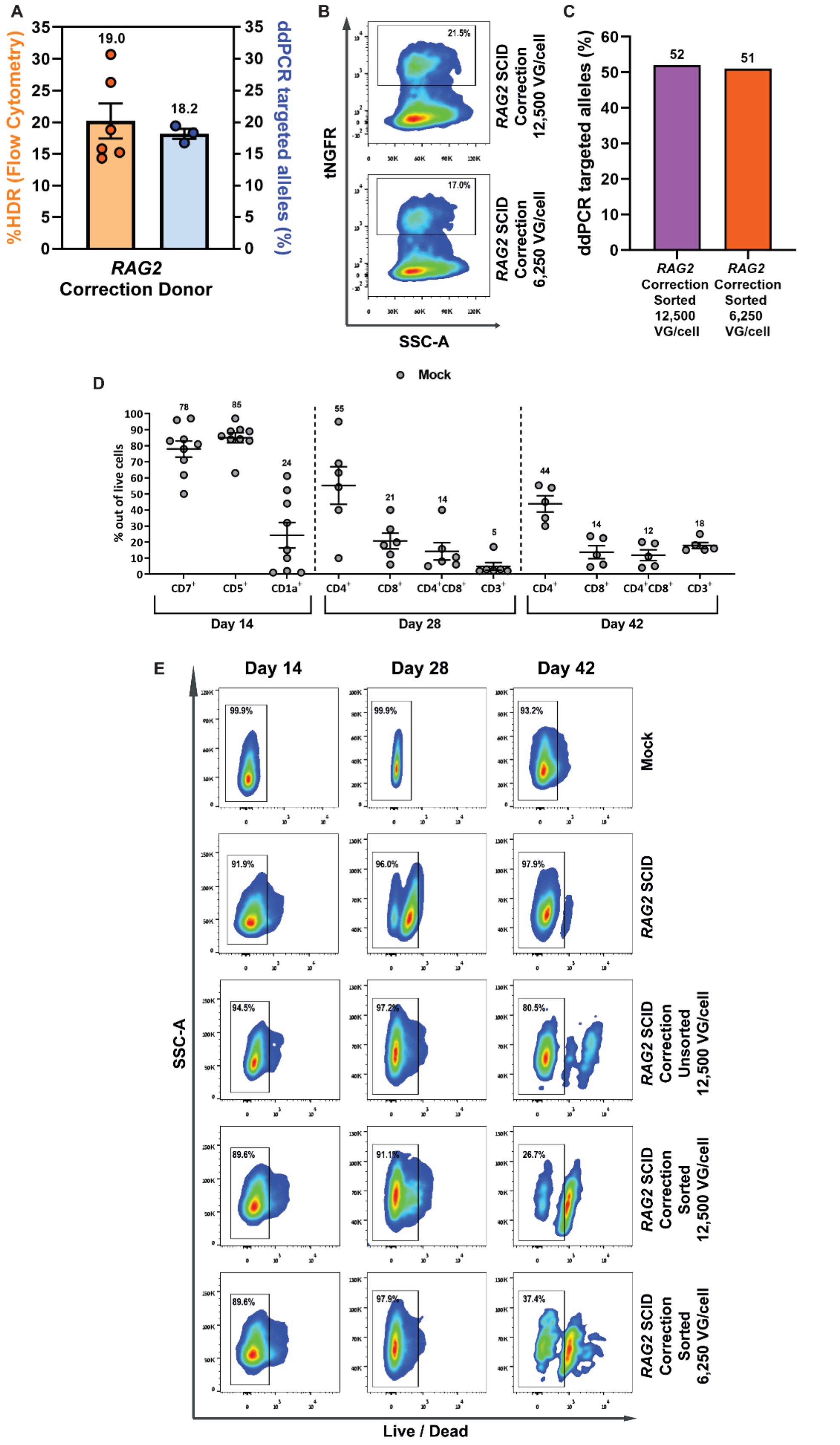
A
%HDR (Flow Cytometry)
ddPCR targeted alleles (%)
19.0
18.2
RAG2
Correction Donor
B
tNGFR
SSC-A
21.5%
17.0%
RAG2 SCID Correction 12,500 VG/cell
RAG2 SCID Correction 6,250 VG/cell
C
ddPCR targeted alleles (%)
52
51
RAG2 Correction Sorted 12,500 VG/cell
RAG2 Correction Sorted 6,250 VG/cell
D
Mock
% out of live cells
78
85
24
55
21
14
5
44
14
12
18
CD7+
CD5+
CD1a+
CD4+
CD8+
CD4+CD8+
CD3+
CD4+
CD8+
CD4+CD8+
CD3+
Day 14
Day 28
Day 42
E
Day 14
Day 28
Day 42
SSC-A
Live / Dead
99.9%
99.9%
93.2%
Mock
91.9%
96.0%
97.9%
RAG2 SCID
94.5%
97.2%
80.5%
RAG2 SCID Correction Unsorted 12,500 VG/cell
89.6%
91.1%
26.7%
RAG2 SCID Correction Sorted 12,500 VG/cell
89.6%
97.9%
37.4%
RAG2 SCID Correction Sorted 6,250 VG/cell

**Figure S6. Diverged *RAG2* cDNA correction in *RAG2*-SCID patient-derived CD34$^+$ HSPCs.** **(A)** HDR frequencies as calculated by flow cytometry and ddPCR for the *RAG2* correction donor tested alone in HD-derived CD34$^+$ HSPCs (Flow cytometry [N=6] and ddPCR [N=3]). **(B)** Representative flow cytometry plots of HDR for corrected *RAG2*-SCID samples. Gating determination is based on cells treated with only the rAAV6 vector to compensate for the episomal expression determined to be around 1%. **(C)** HDR frequencies of the corrected *RAG2*-SCID samples, after sorting, measured by ddPCR and normalized by targeted *CCRL2* alleles. **(D)** Summary of T-cell marker expression of *Mock* cell samples on day 14 (N=9), day 28 (N=6), and day 42 (N=5) of IVTD. **(E)** Representative plots depicting viability staining of samples on days 14, 28, and 42 of IVTD. Gating determination is based on unstained cells. Data are represented as mean ± SEM.

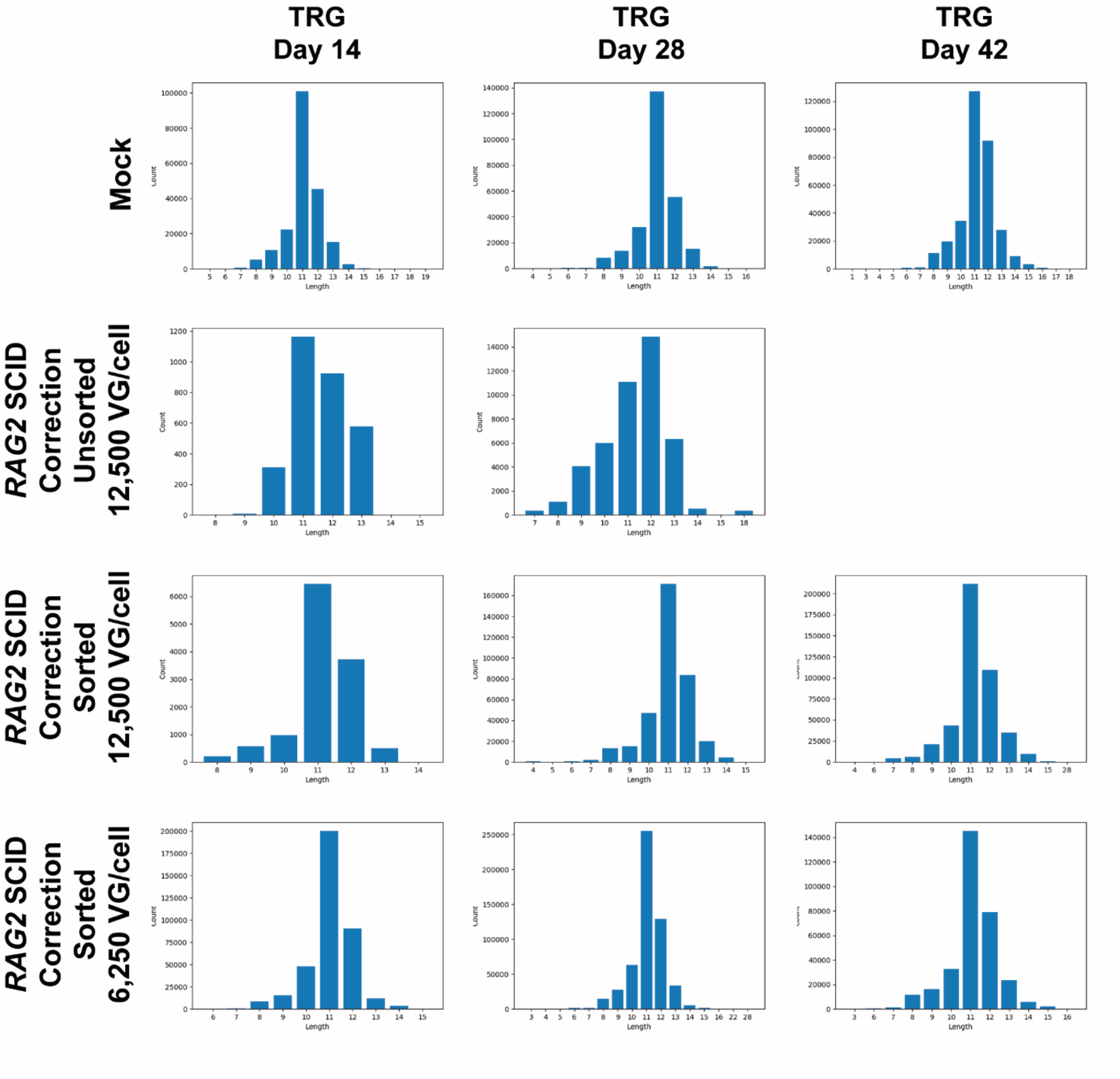


**Figure S7. Diverged *RAG2* cDNA expression in corrected *RAG2*-SCID patient-derived CD34⁺ HSPCs allows for a normal V(D)J recombination process.** Representative CDR3 length distribution among unique TRG sequences expressed in *Mock* and *RAG2*-SCID correction populations on days 14, 28, and 42 of IVTD.

**Table S1. Clinical data of SCID patients**

| | *RAG2*-SCID c.G104T | *RAG1*-SCID c.1407-10 del.4bp-TTGC | |
|---|---|---|---|
| Lymphocyte Subsets | | Before HSCT | After HSCT |
| Lymphocyte Cells (3,400-7,600 cells/$mm^3$) | 1,204 | 2,042 | 1,882 |
| CD3 (2,500-5,500 cells/$mm^3$) | 313 | 0 | 1,618 |
| CD4 (1,600-4,000 cells/$mm^3$) | 229 | 245 | 640 |
| CD8 (600-1,700 cells/$mm^3$) | 193 | 41 | 715 |
| CD20 (300-2,000 cells/$mm^3$) | 0 | 0 | 113 |
| CD56 (% of Total Lymphocytes) (6-30%) | 70 | 91 | 9 |

The cell-subset counts for healthy aged-matched controls are presented in parenthesis[1].

**Table S2. Clinical data of *RAG2*-SCID patient with compound heterozygote mutation p.G95V+E480X.**

Patient presented with Omenn phenotype[2].

| Lymphocyte Subsets | *RAG2*-SCID p.G95V+E480X |
|---|---|
| Lymphocyte Cells (3,400-7,600 cells/$mm^3$) | 10,686 |
| CD3 (2,500-5,500 cells/$mm^3$) | 2,671 |
| CD4 (1,600-4,000 cells/$mm^3$) | 2,351 |
| CD8 (600-1,700 cells/$mm^3$) | 855 |
| CD20 (300-2,000 cells/$mm^3$) | 0 |
| CD56 (% of Total Lymphocytes) (6-30%) | 36 |

The cell-subset counts for healthy aged-matched controls are presented in parenthesis[1].

**Table S3. ddPCR primers and probes**

| | | |
|---|---|---|
| *RAG2* | F | GGGAAGACAATAGCAGG |
| | R | AAGGCATGTATGAGCGT |
| | Probe | FAM/CATCAATAC/ZEN/ATCATCCATGG/3IABkFQ |
| *IL7RA* | F | ACAATAGCAGGCATGCTGG |
| | R | TGAATCCAGTTTGATCTCCTGAC |
| | Probe | FAM/AGTCAGGCA/ZEN/CTGGGTTTGAATGCA/3IABkFQ |
| *DCLRE1C* | F | AACGAAGAATGATTTCTAAGCGCA |
| | R | AATGGCGTTACTGCAGCTAGC |
| | Probe | FAM/TGCTCTGGG/ZEN/AGTTTCGATTT/3IABkFQ |
| *CCR5* | F | GGGAAGACAATAGCAGG |

| | | |
|---|---|---|
| | R | GATGGTGAAGATAAGCCTCAC |
| | Probe | FAM/CTTGTCATG/ZEN/GTCATCTGCTAC/3IABkFQ |
| *CCRL2* | F | GCTGTATGAATCCAGGTCC |
| | R | CCTCCTGGCTGAGAAAAAG |
| | Probe | 5HEX/TGTTTCCTC/ZEN/CAGGATAAGGCAGCTGT/3IABkFQ |

**Table S4. qRT-PCR primers and probes**

| | | |
|---|---|---|
| Endogenous | F | GACATAGTTTCTGATGGTACGTAGA |
| *RAG2* | R | CCAAGTGCTGACAATTAATACCTG |
| | Probe | FAM/TCACGCCTC/ZEN/TCTGAATCTTTGCCG/3IABkFQ |
| Diverged | F | CCCGCTACCTGTACCTTTAAG |
| *RAG2* cDNA | R | TCGCTCACTTCGTTATTGGG |
| | Probe | FAM/TGGTGTTTC/ZEN/TCGCTTTCCAGGGAT/3IABkFQ |
| Endogenous | F | GGTGGATGGCTTAATGCTGAT |
| *DCLRE1C* | R | CAGACCGCAACACTCAGAT |
| | Probe | FAM/TCACGCCTC/ZEN/TCTGAATCTTTGCCG/3IABkFQ |
| *Actin BB* | F | CCTTGCACATGCCGGAG |
| | R | ACAGAGCCTCGCCTTTG |
| | Probe | FAM/TCATCCATG/ZEN/GTGAGCTGGCGG/3IABkFQ |
